# Inhibition of mTOR during a postnatal critical sensitive window rescues deficits in GABAergic PV cell connectivity and social behavior caused by loss of *TSC1*

**DOI:** 10.1101/2020.03.29.014563

**Authors:** Mayukh Choudhury, Clara A. Amegandjin, Vidya Jadhav, Josianne Nunes Carriço, Ariane Quintal, Martin Berryer, Marina Snapyan, Bidisha Chattopadhyaya, Armen Saghatelyan, Graziella Di Cristo

**Author notes:** These authors contributed equally to the study (Mayukh Choudhury, Clara A. Amegandjin). Corresponding author: Graziella Di Cristo, Professor, Department of Neurosciences, Université de Montréal, Centre de Recherche, CHU Sainte-Justine, 3175, Côte-Sainte-Catherine, Montréal, QC H3T 1C5, Canada.

## Abstract

Mutations in regulators of the Mechanistic Target Of Rapamycin Complex 1 (mTORC1), such as *Tsc1/2*, lead to neurodevelopmental disorders associated with autism, intellectual disabilities and epilepsy. Whereas the effects of mTORC1 signaling dysfunction within diverse cell types are likely critical for the onset of the diverse neurological symptoms associated with mutations in mTORC1 regulators, they are not well understood. In particular, the effects of mTORC1 dys-regulation in specific types of inhibitory interneurons are unclear.

Here, we showed that *Tsc1* haploinsufficiency in parvalbumin (PV)-positive GABAergic interneurons either in cortical organotypic cultures or *in vivo* caused a premature increase in their perisomatic innervations, followed by a striking loss in adult mice. This effects were accompanied by alterations of AMPK-dependent autophagy in pre-adolescent but not adult mice. PV cell-restricted *Tsc1* mutant mice showed deficits in social behavior. Treatment with the mTOR inhibitor Rapamycin restricted to the third postnatal week was sufficient to permanently rescue deficits in both PV cell innervation and social behavior in adult conditional haploinsufficient mice. All together, these findings identify a novel role of Tsc1-mTORC1 signaling in the regulation of the developmental time course and maintenance of cortical PV cell connectivity and provide a mechanistic basis for the targeted rescue of autism-related behaviors in disorders associated with deregulated mTORC1 signaling.

## INTRODUCTION

The mechanistic target Of Rapamycin Complex 1 (mTORC1) acts as a central hub integrating internal and external stimuli to regulate many critical cellular processes, including cell growth and metabolism, protein synthesis and autophagy (Saxton and Sabatini, 2017b). mTORC1 signalling has also emerged as an important regulator of brain development and plasticity. De-regulation of mTORC1 signalling network is at the basis of several genetic neurodevelopmental disorders, which share common clinical features, such as epilepsy, autism and other comorbidities (Costa-Mattioli and Monteggia, 2013, Lipton and Sahin, 2014). In particular, mutations in the mTORC1 negative regulators *TSC1* or *TSC2* cause Tuberous Sclerosis Complex (TSC), an autosomal dominant disease associated with high occurrence of epilepsy, intellectual disabilities and autistic traits (Crino et al., 2006). Extensive studies on TSC mutations have set the paradigm for monogenic “mTORpathies”, to understand how mTOR dysregulation affects different processes of brain development (Lipton and Sahin, 2014) and how these may ultimately lead to cognitive and neurological deficits.

The theory of an increased excitation/inhibition (E/I) ratio as an underlying cause of network hyper-excitability and reduced signal-to-noise in the cortex was initially proposed by Rubenstein and Merzenich as a framework for understanding the pathophysiology of autism (Rubenstein and Merzenich, 2003). Over the past 15 years, numerous studies have provided evidence that alterations in E/I balance may be involved in many mouse models of monogenetic autism, however the nature of the underlying mechanisms are heterogeneous thus highlighting that it is critical to understand what sort of circuit alterations are caused by specific genetic mutations (Sohal and Rubenstein, 2019). While numerous studies have focussed on the effects of *Tsc1/2* deletion, and mTOR dysregulation, on cortical and hippocampal excitatory cells (Tavazoie et al., 2005, Bateup et al., 2011, Bateup et al., 2013, Nie et al., 2015), only few studies have addressed whether and how *Tsc1/2* deletion affects cortical GABAergic circuit development (Zhao and Yoshii, 2019, Malik et al., 2019, Artinian et al., 2019, Fu et al., 2012). In particular, whether it plays different roles in specific GABAergic populations is not known.

The neocortex is comprised of a diverse group of inhibitory neurons, which differ in morphology, intrinsic physiological properties and connectivity (Fishell and Rudy, 2011). Among them, Parvalbumin (PV) expressing cells, which represent the largest class of cortical interneurons, specifically target the soma and proximal dendrites of pyramidal cells, and have been implicated in synchronizing the firing of neuronal populations to generate gamma oscillations (Cardin et al., 2009, Sohal et al., 2009, Takada et al., 2014), which in turn allows the cortex to perform precise computational tasks underlying perception, selective attention, working memory and cognitive flexibility in humans and rodents (Fries et al., 2001, Howard et al., 2003, Cho et al., 2006, Fries, 2009). The development of PV cell circuit connectivity is a prolonged process, terminating around the end of adolescence in rodents and primates (Chattopadhyaya et al., 2004, Chattopadhyaya et al., 2007, Baho et al., 2019, Baho and Di Cristo, 2012, Fish et al., 2013). PV cells dysfunction has been found in several mouse models of autism (Selimbeyoglu et al., 2017, Mierau et al., 2016, Chao et al., 2010, Patrizi et al., 2019, Vogt et al., 2018). Conversely, stimulating PV cells has been shown to be sufficient to ameliorate social behaviour (Yizhar et al., 2011, Cao et al., 2018b, Selimbeyoglu et al., 2017). Since mutations in Tsc1 give rise to autistic traits, we questioned whether and how *Tsc1* deletion selectively in PV cells affects their connectivity, and whether and to what extent these alterations in cortical PV cell circuits might be contributing to changes in social behaviour downstream of altered mTOR signaling.

Here, we used a combination of single-cell genetics in cortical organotypic cultures, conditional mutant mice and high-resolution imaging to investigate the effects of TSC-mTORC1 pathway on the development of PV cell connectivity. We found that mutant PV cells (both heterozygous and homozygous) showed a premature increase of their axonal arbor complexity and bouton density in the first three postnatal weeks, followed by a striking loss of connectivity by adulthood. The effect of Tsc1 haploinsufficiency on PV cell connectivity was cell-autonomous and likely mediated by altered AMPK dependent autophagy during a critical postnatal period. Further, conditional mutant mice showed social behaviour deficits. Strikingly, both PV cell connectivity and social behaviour in adult mice were rescued by a short treatment with the mTORC1 inhibitor rapamycin during the third postnatal week, suggesting that inhibiting the premature maturation of PV cell innervations was sufficient to ameliorate the long-term neurological outcomes of the mutation.

## RESULTS

### TSC1 haploinsufficiency in postnatal PV cells reduced PV cell connectivity and altered social behavior in adulthood

The maturation of PV cell innervation is a prolonged process that plateaus at the end of the first postnatal month in mouse cortex (Chattopadhyaya et al., 2004). To investigate whether mTORC1 activation plays a role in this process, we first analyzed the time course of pS6 expression, one of the direct downstream effectors of mTORC1, in PV cells identified by PV immunolabeling (Fig.*1A*). We found that both the proportion of PV cells expressing pS6 (Fig. 1A, B) and the mean intensity of pS6 signal (Fig.1A-C) significantly increased between the second and fourth postnatal weeks in the somatosensory cortex. Similar results were obtained in the visual cortex (data not shown). To investigate whether increase of pS6 expression levels was a generalized phenomenon during this developmental window, we quantified pS6 levels in NeuN+ neurons that represent for the most part pyramidal cells in the cortex (Fig.1D). We found no significant difference in the number of NeuN+ cells expressing pS6 between P18 and P26 (Fig.1E).

**Figure 1.**
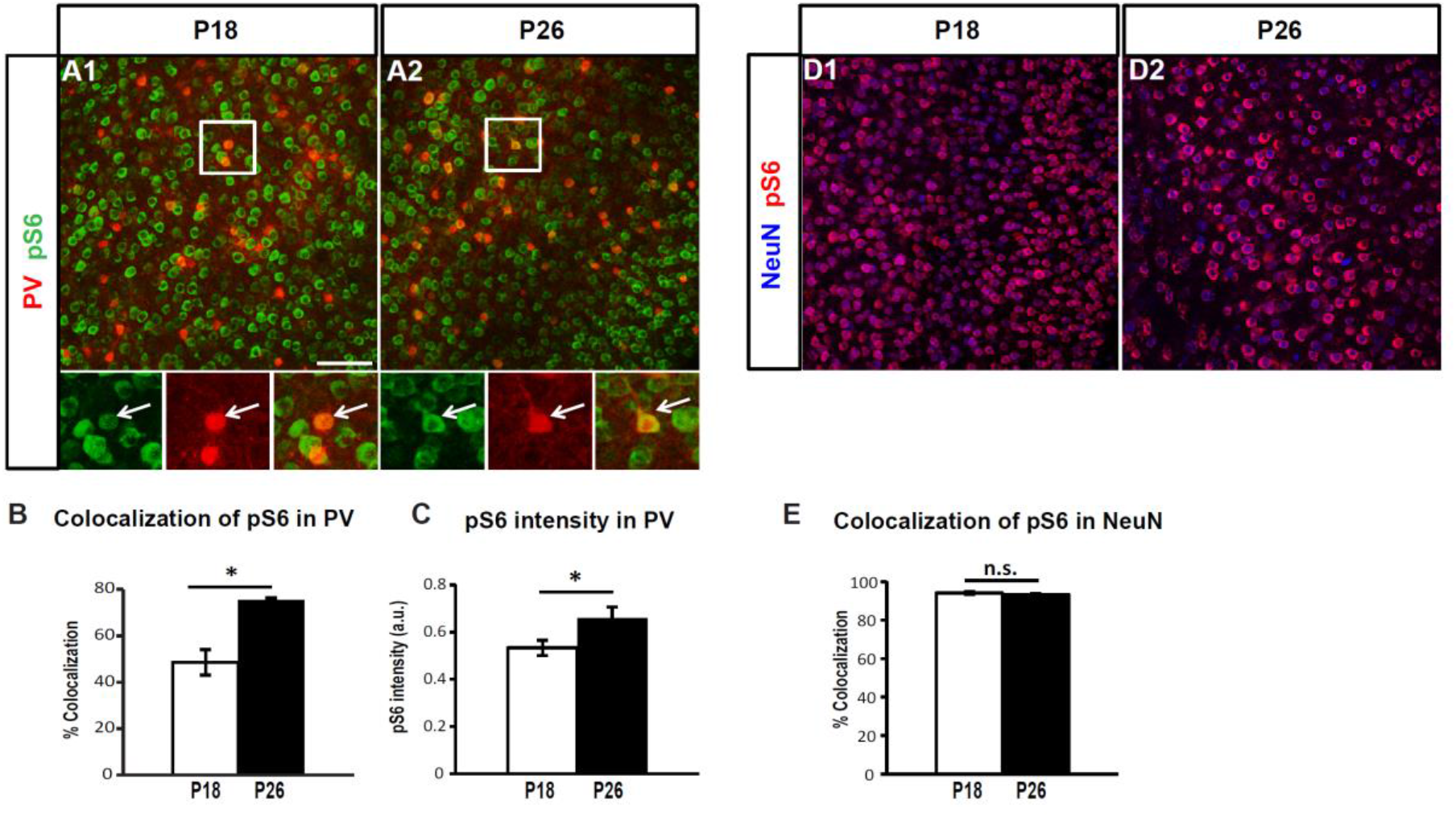
pS6 expression levels increase specifically in PV cells between the second and fourth postnatal weeks. **A**, Coronal sections of mouse somatosensory cortex immunostained for pS6 (green) and PV (red) at P18 (**A1**) and P26 (**A2**). **B**, Number of PV cells expressing detectable levels of pS6 increases during the 2^nd^ to 4^th^ postnatal week (One-Way Anova ***p=0.0005; Tukey’s multiple comparisons test: P14 vs P22 **p= 0.0075; P14 vs P26 **p= 0.0011; P14 vs P33 **p= 0.0042; P18 vs P26 *p= 0.0192). Number of mice; P14, n=5; P18, n=4; P22, n=4; P26, n=3; P33, n=3. **C**, mean pS6 intensity in individual PV cells is significantly higher at P26 than at P18 (Welch’s t-test, **p= 0.0061). Number of mice; P18, n=10; P26, n=7. **D**, Coronal sections of mouse somatosensory cortex immunostained for pS6 (red) and NeuN (blue) at P18 (**D1**) and P26 (**D2**). **E**, Percentage of co-localization of pS6 and NeuN is not significantly different between the two developmental ages (Welch’s t-test, p= 0.7663). Number of mice; P18, n=4; P26, n=3. n.s.: not significant. Scale bars in ***A1-A2, D1-D2***, 75μm. Bar graphs in ***B, C*** and ***E*** represent mean ± SEM.

Since this developmental time window coincides with the peak of the formation of rich and complex perisomatic GABAergic synapse innervation (Chattopadhyaya et al., 2004, Baho et al., 2019), a process that is highly modulated by neuronal activity and sensory experience (Chattopadhyaya et al., 2004, Baho and Di Cristo, 2012), we asked whether and how dysregulation of the TSC-mTOR pathway affects the development of PV cell connectivity.

To answer this question, we used a transgenic mouse carrying a conditional allele of *Tsc1* (Kwiatkowski et al., 2002), which allows cell-specific developmental stage restricted manipulation of *Tsc1*, crossed to the mouse line with the Cre allele under the control of the PV promoter (*PVCre^+/−^*). This cross generated PV-cell restricted homozygous (*PVCre^+/−^; Tsc1^flox/flox^*) and heterozygous (*PVCre^+/−^;Tsc1^flox/+^*) mice and their control *PVCre^−/−^* littermates (*PVCre^−/−^;Tsc1^flox/flox^* and *PVCre^−/−^;Tsc1^flox/+^* mice, referred to hereafter as *Tsc1*^Ctrl^).

To confirm the time course and specificity of Cre expression in *PVCre* mice, we used the *RCE^GFP^* reporter mouse. We observed that about 35% (35±8.11%; n=4 mice) of all PV cells expressed GFP by P14 in the somatosensory cortex, which rose to around 75% (80.43±6.78%; n=8 mice) in P20 mice and to 90% (94.72±2.22%; n=8) in P70 mice, which is consistent with previous findings that PV expression peaks by the 3^rd^ postnatal week. In addition, we confirmed the specificity of Cre expression, since virtually all GFP+ cells expressed PV at all the analyzed ages (98.60±0.50% at P14, 98.21±1.78% at P20; 98.02±0.49% at P70). To control for the efficiency of *Tsc1* deletion in PV cells, we analyzed both pS6 expression and soma size, since both increase following mTORC1 hyperactivity (Tavazoie et al., 2005, Meikle et al., 2007, Tsai et al., 2012). At P45, we observed that a higher proportion of PV cells co-localize with pS6 (Fig.S1 A,B) and that there is a 2.5-fold increase in pS6 intensity in PV cells from *PVCre; Tsc1^flox/flox^* mice (Fig. S1C) as compared to control mice, while the soma size was significantly increased in both mutant genotypes (Fig.S1E).

To determine if *in vivo* postnatal loss of Tsc1 leads to defects in PV cell connectivity, we quantified perisomatic PV synapse density by a) immunostaining cortical slices with pre-synaptic (PV) and post-synaptic (gephyrin) markers and b) by EM analysis of PV+ axons and synapses. We found that the density of perisomatic PV+/gephrin+ punctas was significantly and comparably decreased in mice heterozygous and homozygous for the conditional *Tsc1* allele (Fig.2A-D). In addition, the density of PV+ terminals and the length of PV+ synapses was significantly reduced in *PV-Cre;Tsc1^flox/+^* mice (Fig.2E-L) suggesting that PV-cell restricted, postnatal *Tsc1* haploinsufficiency leads to PV cell hypo-connectivity in adult mice. It has been reported that *Tsc1* deletion in cortical GABAergic neurons (Fu et al., 2012) or Purkinje cells (Tsai et al., 2012) leads to neuronal loss in the targeted population. In our hands, we did not observe any difference in cortical PV cell density in our conditional mutant mice vs control littermates (PV/NeuN; *Tsc1*^Ctrl^ mice: 11.5±1.2%, n=6 mice; *PV-Cre;Tsc1^flox/+^*: 10.4±1.1%, n=4 mice, *PV-Cre;Tsc1^flox/flox^*: 11.6±0.4%, n=5 mice), emphasizing that the observed PV cell hypoconnectivity in adult mutant mice is not due to PV cell loss.

**Figure 2.**
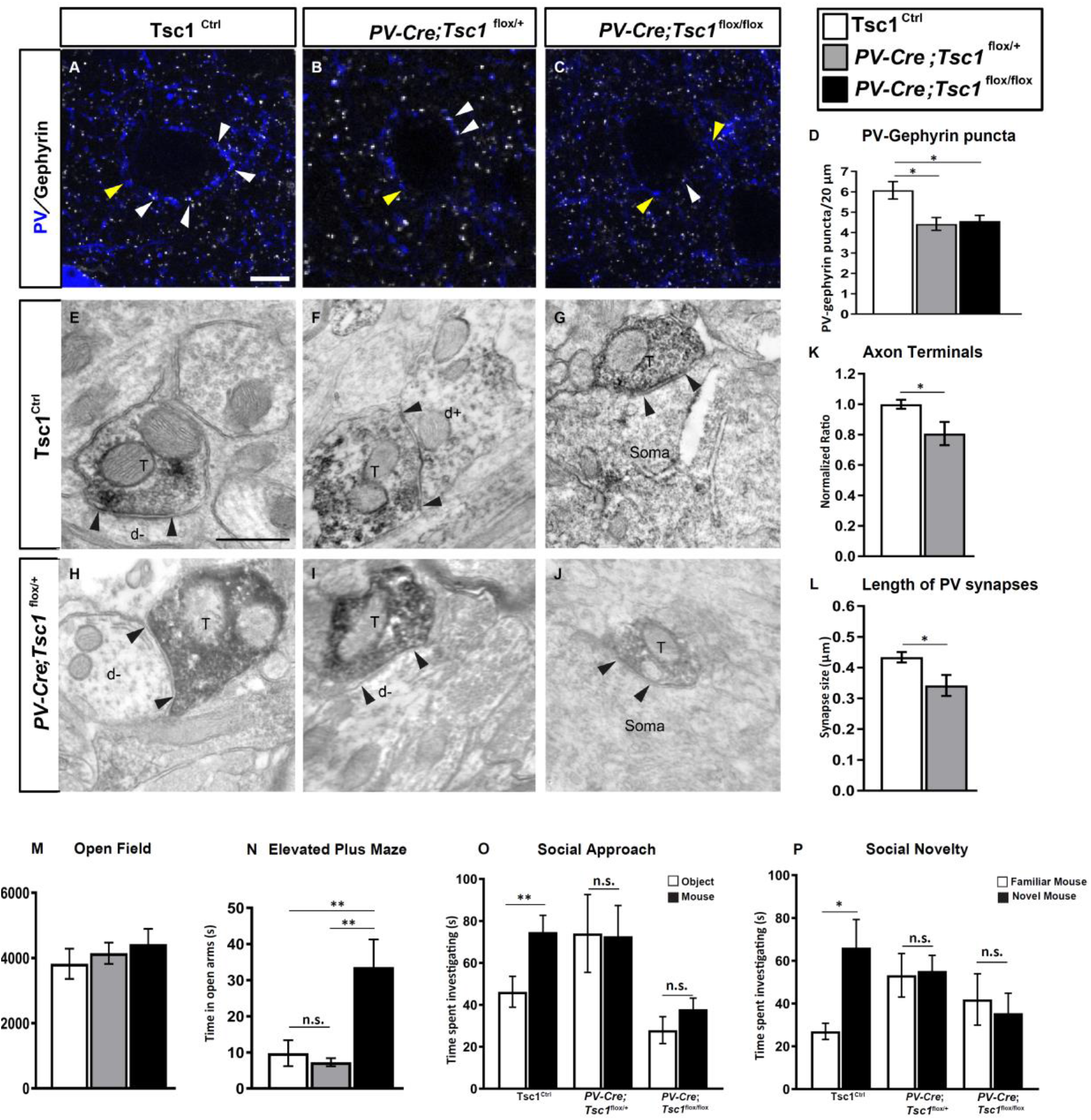
Tsc1 knockout in PV cells causes PV cell hypo-connectivity and social behavioral deficits in young adult mice. **A-C**, Coronal sections of somatosensory cortex immunostained for PV (blue) and gephyrin (grey) in *Tsc1*^Ctrl^ mice at P60 (***A***), *PV-Cre;Tsc1^flox/+^* mice (**B**) and *PV-Cre;Tsc1^flox/flox^* mice (**C**). White arrowheads denote PV-gephyrin colocalized boutons, while yellow arrowheads show PV boutons that do not colocalize with gephyrin puncta. **D**, PV/gephyrin colocalized puncta (one-way ANOVA, *p=0.0096; Tukey’s multiple comparison test: *Tsc1*^Ctrl^ **vs** *PV-Cre;Tsc1^flox/+^* *p=0.0140; *Tsc1*^Ctrl^ **vs** *PV-Cre;Tsc1^flox/flox^* *p=0.0236), n=5 mice for all genotypes. Scale bar: 10 μm. **E-J,** PV-immunolabeled axon terminals in somatosensory cortex of *Tsc1*^Ctrl^ (**E, F, G**) and *PV-Cre;Tsc1^flox/+^* (**H, I, J**) at P60. **E-G**, Labeled axon terminals (T) make symmetric synaptic contact (flanked by arrowheads) with an unlabeled dendritic shaft (d-). **F**, Rare symmetric synapse between PV positive axon terminal and labeled dendrite (d+). **G** and **J:** synaptic contact between a PV+ axon terminal (T) and an unlabeled cell soma. **K**, Quantification of PV+ axon terminals in *PV-Cre;Tsc1^flox/+^* mice (Unpaired t-test, *p=0.0453). **L**, *PV-Cre; Tsc1^flox/+^* mice have shorter synapses than *Tsc1*^Ctrl^ (Unpaired t test, *p=0.0464). Number of mice: *Tsc1*^Ctrl^, n=4; *PV-Cre;Tsc1^flox/+^* mice, n=3. Scale bars: **E-J**, 500 nm. **M**, Open field test: Quantification of distance travelled during exploratory activity in an open field arena at P33 shows that distances covered in *PV-Cre;Tsc1^flox/flox^* and *PV-Cre;Tsc1^flox/+^* mice are similar to *Tsc1*^Ctrl^ (one-way ANOVA, p>0.05; Tukey’s multiple comparisons test: *Tsc1*^Ctrl^ **vs** *PV-Cre;Tsc1^flox/+^* p=0.8537; *Tsc1*^Ctrl^ **vs** *PV-Cre;Tsc1^flox/flox^* p=0.5794; *PV-Cre;Tsc1^flox/+^* **vs** *PV-Cre;Tsc1^flox/flox^* p=0.8956). Number of mice: *Tsc1*^Ctrl^, n=12; *PV-Cre;Tsc1^flox/+^* mice, n=10; *PV-Cre;Tsc1^flox/flox^*, n= 10. **N**, Elevated plus maze: Quantification of time spent in the open arms of elevated plus maze arena at P35 shows reduced anxiety-like behaviour in *PV-Cre;Tsc1^flox/flox^* mice (one-way ANOVA, ***p=0.0006; Tukey’s multiple comparisons test: *Tsc1*^Ctrl^ **vs** *PV-Cre;Tsc1^flox/+^* p=0.9205; *Tsc1*^Ctrl^ **vs** *PV-Cre;Tsc1^flox/flox^* **p=0.0017; *PV-Cre;Tsc1^flox/+^* **vs** *PV-Cre;Tsc1^flox/flox^* **p=0.0018). Number of mice: *Tsc1*^Ctrl^, n=21; *PV-Cre;Tsc1^flox/+^* mice, n=13; *PV-Cre;Tsc1^flox/flox^* n=13. **O, P**, Unlike *Tsc1*^Ctrl^ mice, both *PV-Cre;Tsc1^flox/+^* and *PV-Cre;Tsc1^flox/flox^* mice failed to show preference for a mouse vs an object (**O**) or for a novel mouse vs a familiar one (**P**) (two-way ANOVA with Tukey’s post hoc analysis, *p<0.05, **p<0.01); Number of mice: (**O**) *Tsc1*^Ctrl^, n = 15; *PV-Cre;Tsc1^flox/+^*. n=11; *PV-Cre;Tsc1^flox/flox^*, n=11. (**P**) *Tsc1*^Ctrl^, n=11; *PV-Cre;Tsc1^flox/+^*, n=14; *PV-Cre;Tsc1^flox/flox^*, n= 12. Data represent mean ± SEM.

To investigate whether *PV-Cre; Tsc1^flox^* mutants might show abnormal behaviors resembling ASDs, we evaluated social interaction using the three-chamber assay of social approach and preference for social novelty. Both heterozygous and homozygous mutant mice showed no significant differences in the time spent interacting with a mouse versus an object (Fig.2O) or with a novel versus a familiar mouse (Fig.2P). Further, there were no differences in locomotor activity, as tested in the open field (Fig.2M), and no increased anxiety in the elevated plus maze paradigm (Fig.2N). In fact, *PV-Cre;Tsc1^flox/flox^* but not *PV-Cre;Tsc1^flox/+^* mice exhibit less anxiety-like behavior, since they spent significantly more time in the open arms (Fig.2N).

Since *Tsc1* deletion has been shown to affect glutamatergic synapse formation (Tavazoie et al., 2005, Bateup et al., 2011), we analyzed the density of asymmetrical synapses (which represent glutamatergic inputs) onto PV cell dendrites by EM (Fig.S2A-D). We found that while the density of PV-positive dendrites was not significantly different between *PV-Cre;Tsc1^flox/+^* mice and wild-types littermates (Fig.S2E), the density of dendrites bearing asymmetrical synapses was significantly reduced in the mutant mice (Fig.S2F), suggesting that Tsc1-mediated signaling promotes the formation of excitatory inputs onto PV cells.

In summary, *Tsc1* deletion in postnatal PV cells leads to alterations in both PV cell connectivity and in social behavior. PV cell hypo-connectivity in adult mutant mice could be directly caused by Tsc1-mTORC1 signaling dysregulation in PV cells or induced as a consequence of homeostatic feedback mechanisms that influence neural circuit development. We then used both *in vitro* and *in vivo* approaches to determine the cell-autonomous and network phenotypes resulting from the genetic deletion of *Tsc1* in cortical PV cells.

### mTORC1 hyperactivation in single PV cells induced a premature increase in bouton density and axon branching, subsequently followed by excessive bouton pruning

To explore the cell autonomous effects of *Tsc1* deletion or haploinsufficency during specific developmental phases of PV cell connectivity, we used single cell genetic manipulation in cortical organotypic cultures (Fig.S3A). To reduce *Tsc1* expression in single PV cells and simultaneously labelling their axons and synapses, we used a previously characterized promoter region P_G67_ (Chattopadhyaya et al., 2004) to express either Cre recombinase together with GFP (P_G67_-GFP/Cre) or GFP alone (control) in single PV cells in cortical organotypic cultures from *Tsc1^flox/flox^* and *Tsc1^flox/+^* mice. This approach allowed us to generate specifically *Tsc1*^−/−^ and *Tsc1*^+/−^ PV cells in an otherwise wild-type background. Deletions of either one or both *Tsc1* alleles significantly increased pS6 expression levels in the transfected sparse PV cells (Fig.S3E), while cell soma size was significantly increased only in *Tsc1^−/−^* PV cells (Fig. S3F).

We have previously shown that the basic features of the mature perisomatic innervation formed by PV cells onto pyramidal cells are recapitulated in cortical organotypic cultures (Di Cristo et al., 2004, Chattopadhyaya et al., 2004). PV innervation starts out with simple axons, which develop into complex, highly branched arbors in the subsequent 4 weeks with a time course similar to that observed *in vivo* (Chattopadhyaya et al., 2004). In particular, PV cell axonal arborization and bouton density increase significantly between EP18 (P5+13 days *in vitro=Equivalent Postnatal day 18)* and EP24. To investigate the effect of premature mTORC1 activation on PV cell synapse innervation, we biolistically transfected PV cells at EP10 and analyzed them at EP18 (Fig.*S3A*). Following *Tsc1* deletion, we quantified two aspects of individual PV cell connectivity – 1) the extent of perisomatic innervation around single targeted somata (terminal branching and perisomatic bouton density) and 2) the fraction of potentially innervated somata within the basket cell arbor (percentage of innervation or the innervation field). We have previously shown that the vast majority of GFP-labeled boutons in our experimental conditions most likely represent presynaptic terminals (Chattopadhyaya et al., 2004, Wu et al., 2012, Chattopadhyaya et al., 2007). We found that both *Tsc1*^−/−^ and *Tsc1*^+/−^ PV cells formed premature perisomatic innervations, characterized by increased bouton density (Fig.3A, B, E, Fig. S4A) and terminal axonal branching around NeuN+ contacted somata (Fig.3F, Fig.S4B), and increased percentage of contacted target cells (Fig.3H). To determine whether the effects of *Tsc1* deletion are due to mTORC1 hyperactivation, we treated cortical organotypic cultures with the mTORC1 inhibitor Rapamycin from EP10-18 (90ng/ml, Fig.S5) and found that Rapamycin treatment reversed the increase in bouton density in *Tsc1^−/−^* PV cells (Fig.S5E) as well as terminal branching (Fig. S5F). All together, these data suggest that mTORC1 hyper activation leads to the premature formation of PV cell innervations in a cell-autonomous manner.

**Figure 3:**
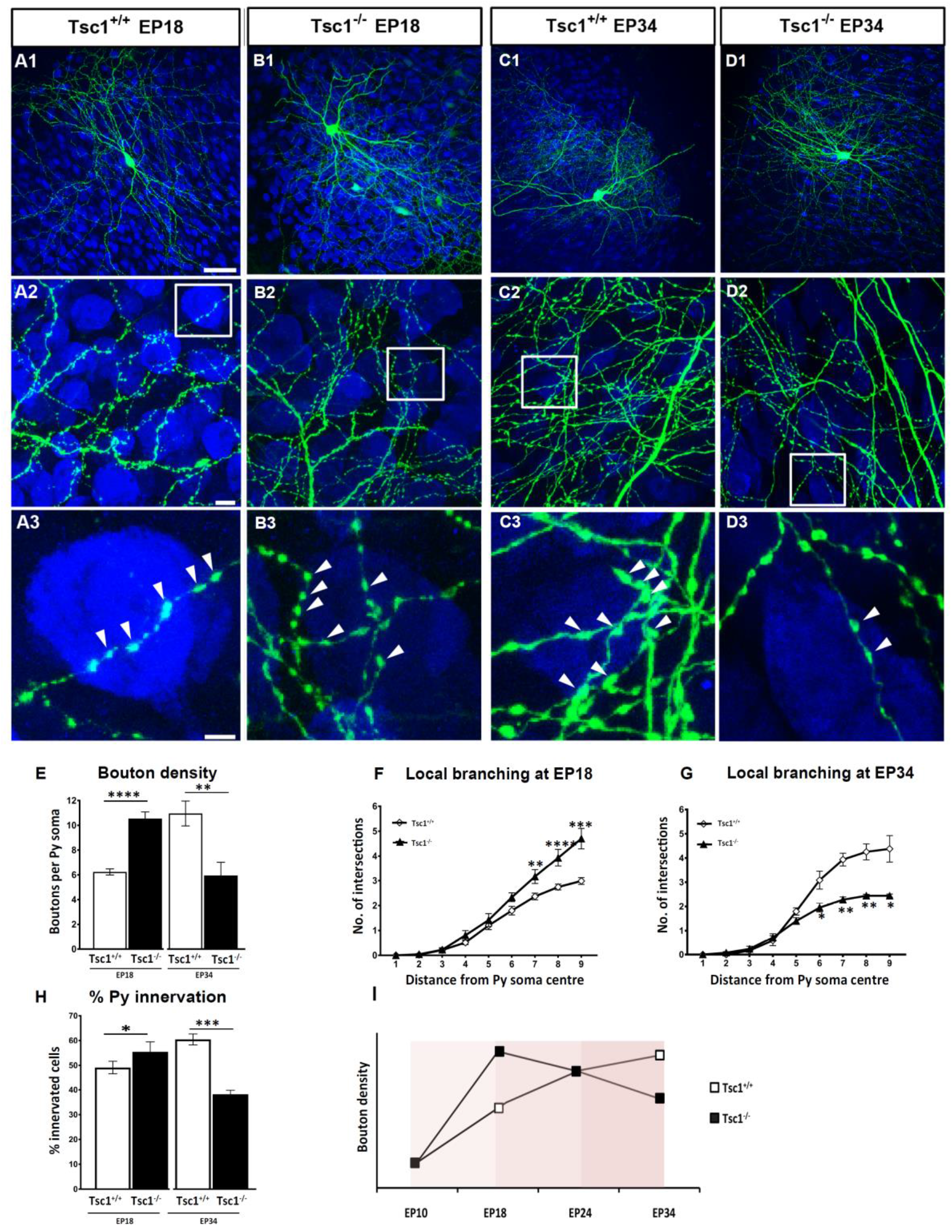
*Tsc1* knockout in single PV neurons causes a premature increase in axonal terminal branching and bouton density followed by excessive pruning. **A1**, EP18 *Tsc1^+/+^* PV cell showing characteristic branching (**A2**) and boutons (**A3**, arrowheads) on the postsynaptic somata identified by NeuN immunostaining (blue). **B**, *Tsc1*^−/−^ PV cells lacking both alleles (**B1-B3**) of *Tsc1* show significant increase in bouton density at EP18. **C, Control** EP34 *Tsc1^+/+^* PV cell. (**D1-D3**), EP34 *Tsc1^−/−^* PV cell showing significantly decreased axonal branching (**C2 vs D2**) and perisomatic boutons (**C3 vs D3**). **E**, *Tsc1^−/−^* PV cells show an increase in bouton density at EP18 (Welch’s t-test, ****p<0.0001) followed by a decrease at EP34 (Welch’s t-test, **p=0.0091). **F**, *Tsc1^−/−^* PV cells show more developed branching than *Tsc1^+/+^* cells at EP18 (Welch’s t-test:**p=0.0017 (radius7), ****p<0.0001 (radius8), ***p=0.0001 (radius9)). **G**, At EP34 *Tsc1^−/−^* PV cells are characterized by simpler axonal branching compared to controls (Welch’s t-test: *p=0.0336 (radius6), **p=0.0016 (radius7), **p=0.0047 (radius8), *p=0.0232 (radius9)). **H**, Percentage of innervation is significantly increased in *Tsc1^−/−^* PV cells at EP18 and reduced at EP34 (Welch’s t-test: EP18,*p=0.0114; EP34, ****p<0.0001). Number of PV cells: At EP18; n=16 for *Tsc1^+/+^*, n=9 for *Tsc1^−/−^*. At EP34: n=5 *Tsc1^+/+^*, n=6 *Tsc1^−/−^*. **I**, Schematic representation of bouton density during the postnatal maturation of *Tsc1^+/+^* and *Tsc1^−/−^* PV cells Scale bars: ***A1-D1***, 50 μm; ***A2-D2***, 10 μm; ***A3-D3***, 5 μm. Data represents mean ± SEM.

Next, we asked whether the premature development of PV cell innervation was long lasting. As described above, PV cells were transfected at EP10 and then analyzed either at EP24 (Fig. S6, during the peak of the proliferation of PV cell innervation) or at EP34 (Fig.3 C, D, when PV cell innervation has matured and is stable). At EP24, perisomatic innervation formed by *Tsc1^−/−^* PV cells were similar to those formed by age-matched wild-type cells (Fig.S6). However, at EP34, *Tsc1^−/−^* PV cells showed significantly poorer innervations than age-matched *Tsc1^+/+^* PV cells (Fig.3E, G, H). All together, these data show that dysregulated TSC-mTORC1 signaling in individual PV cells alters the development of their innervations, inducing first a premature increase in axonal branching and bouton density followed by excessive pruning (Fig. 3I), in a cell-autonomous fashion.

Next, we sought to investigate whether the phenotypic switch of PV cell connectivity caused by *Tsc1* deletion occurs *in vivo.* Since Cre expression in *PVCre* mice starts at around P10 and only peaks towards the third postnatal week, we reasoned that the time course of *Tsc1* allele recombination, and its effects on PV cell innervation, may be highly variable between P18 and P24. To overcome this issue, we needed a specific early promoter expressing Cre to knockdown Tsc1 in PV neurons and therefore used the Tg(*Nkx2.1-Cre*) for this purpose. We generated Tg(*Nkx2.1-Cre*);*Tsc1^flox^* and control littermates. NKX2.1 is a transcription factor expressed at E10.5 by GABAergic cell precursors in the medial ganglionic eminence (MGE), which gives rise to cortical PV- and somatostatin-expressing (SST) GABAergic cells (Xu et al., 2008). At P18, pS6 levels and soma size were significantly increased in PV cells from Tg(*Nkx2.1-Cre*);*Tsc1^flox/flox^* mouse somatosensory cortex (Fig.S7A-C, E). By P45, PV cells showed a four and two-fold increase in pS6 intensity in Tg(*Nkx2.1-Cre*); *Tsc1^flox/flox^* and Tg(*Nkx2.1-Cre*);*Tsc1^flox/+^* mice compared to control mice, respectively (Fig. S7D). PV cell somata were larger in Tg(*Nkx2.1-Cre*);*Tsc1^flox/+^* compared to *Tsc1*^Ctrl^ mice, even if not as large as those in Tg(*Nkx2.1-Cre*); *Tsc1^flox/flox^* mice (Fig.S7F), suggesting that deletion of one *Tsc1* allele may have slow, cumulative effects *in vivo*, consistent to what is previously reported in Purkinje cell-specific *Tsc1* mutant mice (Tsai *et al.*, 2012). A previous study showed that conditional *Tsc1* deletion in GABAergic progenitor cells using Dlx5/6-Cre mice leads to reduced cortical GABAergic cell density (Fu et al., 2012). Conversely, we found no difference in PV cell density between the mutant mice and control littermates at P18 (PV/NeuN; Ctrl mice: 8.9±0.7%; n=3 mice; *Nkx2.1-Cre;Tsc1^flox/+^*: 8.1±0.3%; n=3 mice, *Nkx2.1-Cre;Tsc1^flox/flox^*: 7.9±0.3%; n=4 mice). This difference suggest that conditional deletion of *Tsc1* at the time of cell cycle exit (Nkx2.1-Cre) has a different impact than removal on the mantle zone (Dlx5/6-Cre) on GABAergic neuron survival.

In order to analyze the PV cell axonal morphology at high resolution we turned to organotypic cultures from Tg(*Nkx2.1-Cre*); *Tsc1^flox/flox^*, Tg(*Nkx2.1-Cre*)*Tsc1^flox/+^* and *Tsc1*^Ctrl^ mice transfected with P_G67_-GFP at different developmental stages (Fig. 4, 5). At EP18, before the peak of PV cell synapse proliferation, similar to what we observed with the single cell *Tsc1* deletion, we found that PV cells from both Tg(*Nkx2.1-Cre*)*Tsc1^flox/flox^* and Tg(*Nkx2.1-Cre*);*Tsc1^flox/+^* mice formed more complex perisomatic innervations, characterized by increased perisomatic bouton density (Fig.4A-D) and terminal branching (Fig.4E) compared to age-matched control PV cells in cultures transfected from *Tsc1*^Ctrl^ mice.

**Figure 4.**
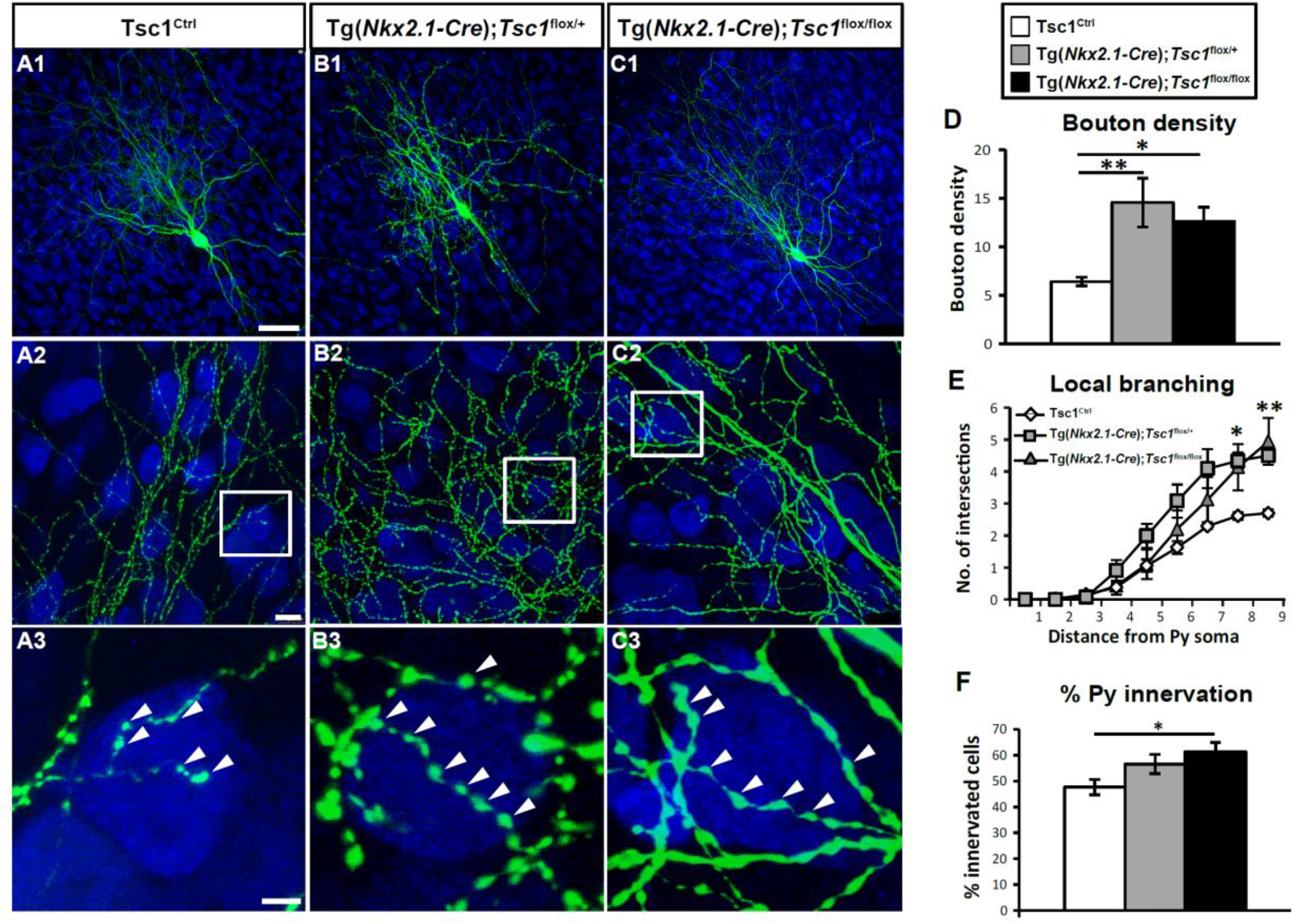
PV cells show prematurely rich perisomatic innervation in Tg(*Nkx2.1-Cre*);*Tsc1^flox/flox^* and Tg(*Nkx2.1-Cre*);*Tsc1^flox/+^* mice at EP18. **A1**, A PV cell (green) amongst NeuN immunostained neurons (in blue) in cortical organotypic culture from a *Tsc1*^Ctrl^ mouse at EP18. **A2**, PV cell from *Tsc1*^Ctrl^ slice shows characteristic branching and multiple boutons (arrowheads) on the postsynaptic somata (**A3**). PV cells from Tg(*Nkx2.1-Cre*);*Tsc1^flox/+^* mice (**B1-B3**) and Tg(*Nkx2.1-Cre*);*Tsc1^flox/flox^* mice (**C1-C3**) show increased bouton density (**D**) (one-way ANOVA, **p=0.0039; Holm-Sidak post hoc analysis: *Tsc1*^Ctrl^ **vs** Tg(*Nkx2.1-Cre*);*Tsc1^flox/+^* **p=0,0023; *Tsc1*^Ctrl^ vs Tg(*Nkx2.1-Cre*);*Tsc1^flox/flox^* *p=0.0242). Number of mice: *Tsc1*^Ctrl^ n=7, Tg(*Nkx2.1-Cre*);*Tsc1^flox/+^* n=7, Tg(*Nkx2.1-Cre*); *Tsc1^flox/flox^* n=6. ***E***, Local branching (one-way ANOVA *p=0.0113 (Radius 8), **p=0.0096 (Radius 9); Holm-Sidak post hoc analysis: (Radius 8) *Tsc1*^Ctrl^ **vs** Tg(*Nkx2.1-Cre*); *Tsc1^flox/+^* *p=0,0155; *Tsc1*^Ctrl^ **vs** Tg(*Nkx2.1-Cre*); *Tsc1^flox/flox^* *p=0.0425, Tg(*Nkx2.1-Cre*); *Tsc1^flox/+^* **vs** Tg(*Nkx2.1-Cre*); *Tsc1^flox/flox^* p=0.8062; (Radius 9) *Tsc1* **vs** Tg(*Nkx2.1-Cre*); *Tsc1^flox/+^* *p=0.0317; *Tsc1*^Ctrl^ **vs** Tg(*Nkx2.1-Cre*); *Tsc1^flox/flox^* *p=0.0148, Tg(*Nkx2.1-Cre*); *Tsc1^flox/+^* **vs** Tg(*Nkx2.1-Cre*); *Tsc1^flox/flox^* p=0.9738). Number of mice: *Tsc1*^Ctrl^ n=7, Tg(*Nkx2.1-Cre*);*Tsc1^flox/+^* n=5, Tg(*Nkx2.1-Cre*); *Tsc1^flox/flox^* n=6. **F**, Percentage of innervation (one-way ANOVA, *p=0.0254; Holm-Sidak post hoc analysis: *Tsc1*^Ctrl^ **vs** Tg(*Nkx2.1-Cre*); *Tsc1^flox/+^* p=0,0823; *Tsc1*^Ctrl^ **vs** Tg(*Nkx2.1-Cre*); *Tsc1^flox/flox^* *p=0.0168). Number of mice: *Tsc1*^Ctrl^ n=8, Tg(*Nkx2.1-Cre*); *Tsc1*^Ctrl^ n=6, Tg(*Nkx2.1-Cre*); *Tsc1^flox/flox^* n=7. Scale bars: ***A1-C1***, 20 μm; ***A2-C2*** 10 μm, ***A3-C3***, 3 μm. Data represents mean ± SEM.

Conversely, at EP34, in a period when PV axonal arbor maturation has reached stability, PV neurons from both genotypes (Tg(*Nkx2.1-Cre*);*Tsc1^flox/flox^*, Tg(*Nkx2.1-Cre*);*Tsc1^flox/+^*) showed significantly reduced perisomatic bouton density (Fig.5A-C, D), terminal branching (Fig.5E) and innervated a smaller percentage of pyramidal neurons (Fig.5F). Overall, these results confirm that embryonic deletion of *Tsc1* has opposite effects on PV perisomatic synapse formation and maintenance, initially accelerating the formation of PV synaptic innervation and subsequently impairing perisomatic synapses at the maturation phase. Further, deletion of a single *Tsc1* allele in PV cells is sufficient to alter its connectivity both at the single cell and network levels.

**Figure 5.**
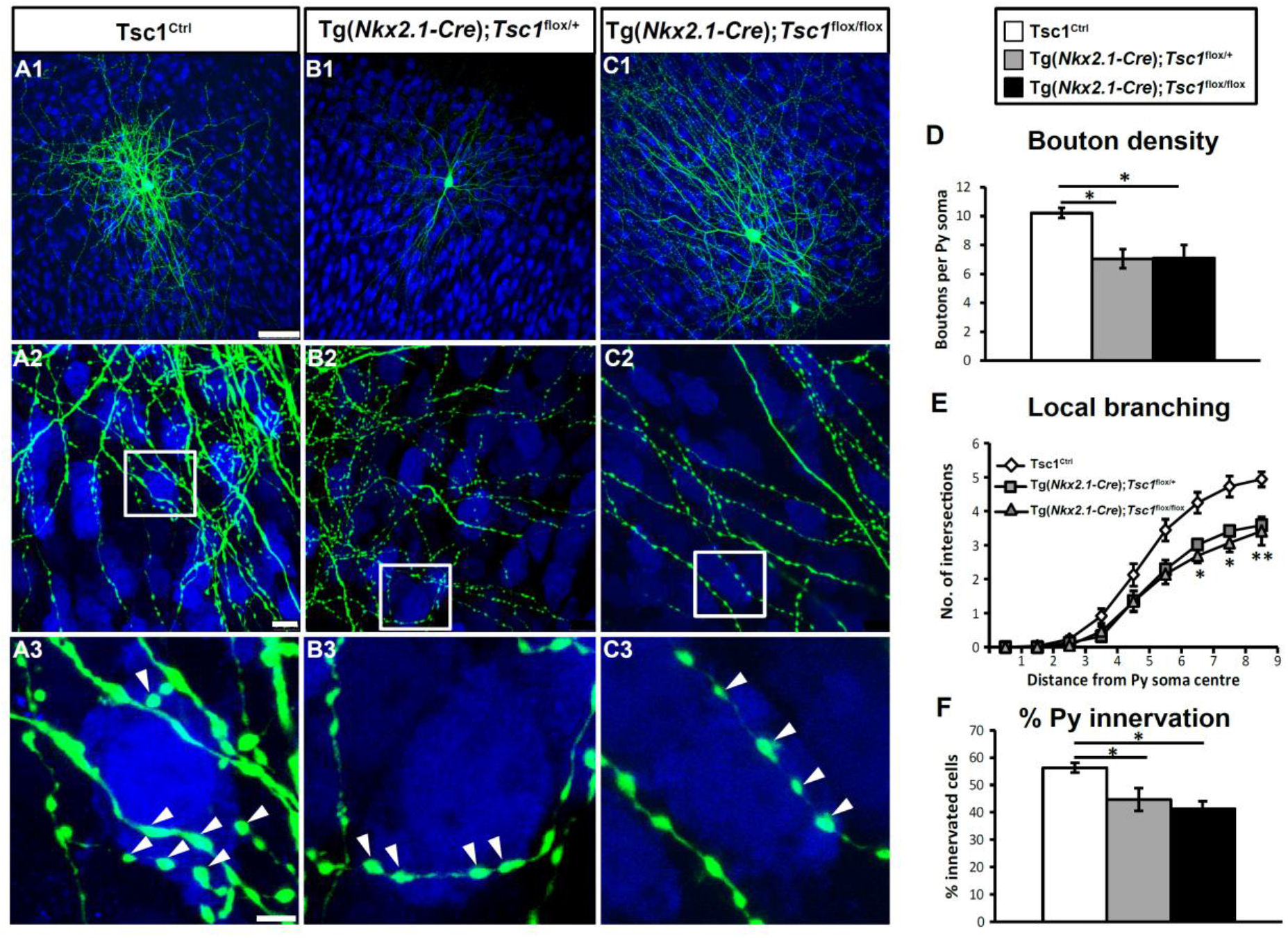
PV interneurons show significantly reduced perisomatic innervation in Tg(*Nkx2.1-Cre*);*Tsc1^flox/^flox^^* and Tg(*Nkx2.1-Cre*);*Tsc1^flox/+^* mice at EP34. **A**, A PV cell (green) among NeuN immunostained neurons (blue) in cortical organotypic cultures from a Tsc1^Ctrl^ mouse at EP34. **B, C,** PV cells from Tg(*Nkx2.1-Cre*);*Tsc1^flox/+^* mice (**B1-B3**) or Tg(*Nkx2.1-Cre*);*Tsc1^flox/flox^* mice (**C1-C3**) show decreased bouton density (**D**) (one-way ANOVA, *p=0.0157; Holm-Sidak post hoc analysis: *Tsc1*^Ctrl^ **vs** Tg(*Nkx2.1-Cre*);*Tsc1^flox/+^ *p=0,0214; *Tsc1*^Ctrl^ **vs** Tg(*Nkx2.1-Cre*);*Tsc1^flox/flox^* *p=0.0214). Local branching (**E**) (one-way ANOVA, **p=0.0040 (Radius 7), *p=0.0127 (Radius 8), **p=0.0011 (Radius 9); Holm-Sidak post hoc analysis: (Radius 7)*Tsc1* ^Ctrl^ **vs** Tg(*Nkx2.1-Cre*);Tsc1^flox/+^* **p=0,0032; *Tsc1*^Ctrl^ **vs** Tg(*Nkx2.1-Cre*);*Tsc1^flox/flox^* *p=0.0117; (Radius 8) *Tsc1*^Ctrl^ **vs** Tg(*Nkx2.1-Cre*); *Tsc1^flox/+^* *p=0,0126; *Tsc1^Ctrl^* **vs** Tg(*Nkx2.1-Cre*);*Tsc1^flox/flox^* *p=0.0149; (Radius 9) *Tsc1*^Ctrl^ **vs** Tg(*Nkx2.1-Cre*); *Tsc1^flox/+^* ***p=0,0008; *Tsc1*^Ctrl^ **vs** Tg(*Nkx2.1-Cre*);*Tsc1^flox/flox^* **p=0.0034). Number of mice: *Tsc1*^Ctrl^ n=6, Tg(*Nkx2.1-Cre*);*Tsc1^flx/+^* n=6, Tg(*Nkx2.1-Cre*);*Tsc1^flox/+^* n=6. **F**, Percentage of innervation is also significantly lower for PV cells from Tg(*Nkx2.1-Cre*);*Tsc1^flox/+^* and Tg(*Nkx2.1-Cre*);*Tsc1^flox/flox^* mice (one-way ANOVA, *p=0.0212, Holm-Sidak post hoc analysis: *Tsc1*^Ctrl^ **vs** Tg(*Nkx2.1-Cre*);*Tsc1^flox/+^* *p=0,0257; *Tsc1*^Ctrl^ **vs** Tg(*Nkx2.1-Cre*);*Tsc1^flox/flox^* *p=0.0179). Number of mice: *Tsc1*^Ctrl^ n=6, Tg(*Nkx2.1-Cre*);*Tsc1^flox/+^* n=8, Tg(*Nkx2.1-Cre*); *Tsc1^flox/flox^* n=6. Arrowheads indicate boutons. Scale bars: ***A1-C1***, 20 μm; ***A2-C2*** 10 μm, ***A3-C3***, 3 μm. Data represents mean ± SEM.

Furthermore, behavioral analysis showed that both heterozygous and homozygous Tg(*Nkx2.1-Cre*);*Tsc1^flox^* mice phenocopied the deficits in social behavior (social approach and social novelty preference) that we found in *PV-Cre;Tsc1^flox^* mice (Fig. S8).

### Tsc1 deletion in GABAergic cells causes transient autophagy dysfunction in adolescent mice

A recent study revealed that, in neurons, Tsc2/1 loss altered the process of autophagy by AMP-activated kinase (AMPK)-dependent mechanisms (Di Nardo et al., 2014). To investigate whether autophagy was affected by conditional *Tsc1* deletion, we analyzed by western blot the expression levels of the autophagosomal lipidated microtubule-associated protein 1 light chain 2 (LC3-II) and of the autophagy substrate p62/sequestosome 1 in *Nkx2.1Cre;Tsc1^flox/flox^* mice compared to that of their control littermates at P14, when we observed premature formation of PV cell synapses (Fig.4), and P40, when PV cells displayed significantly reduced connectivity (Fig.5) by western blot. Since GABAergic neurons constitute a minority of cortical cells, we extracted proteins from the olfactory bulb where GABAergic cells are highly enriched, to increase the likelihood of detecting small changes in protein expression levels. We found significantly increased levels of LC3-II, but not of p62, in P14 mutant mice compared to their littermates (Fig.6 A,B). We further found increased activation of AMPK, as indicated by the expression levels of phospho-AMPK (at T172) (Fig.6 C,D), consistent with what was previously reported (Di Nardo et al., 2014). P40 mice, on the other hand, did not show any significant alterations in any of these markers (Fig.6E-H). Overall, these data suggest that *Tsc1* loss in MGE-derived GABAergic cells leads to a dys-regulation of autophagy and of AMPK activation during a critical developmental window, which overlaps temporally with the maturation of PV cell connectivity (Chattopadhyaya et al., 2004).

**Figure 6.**
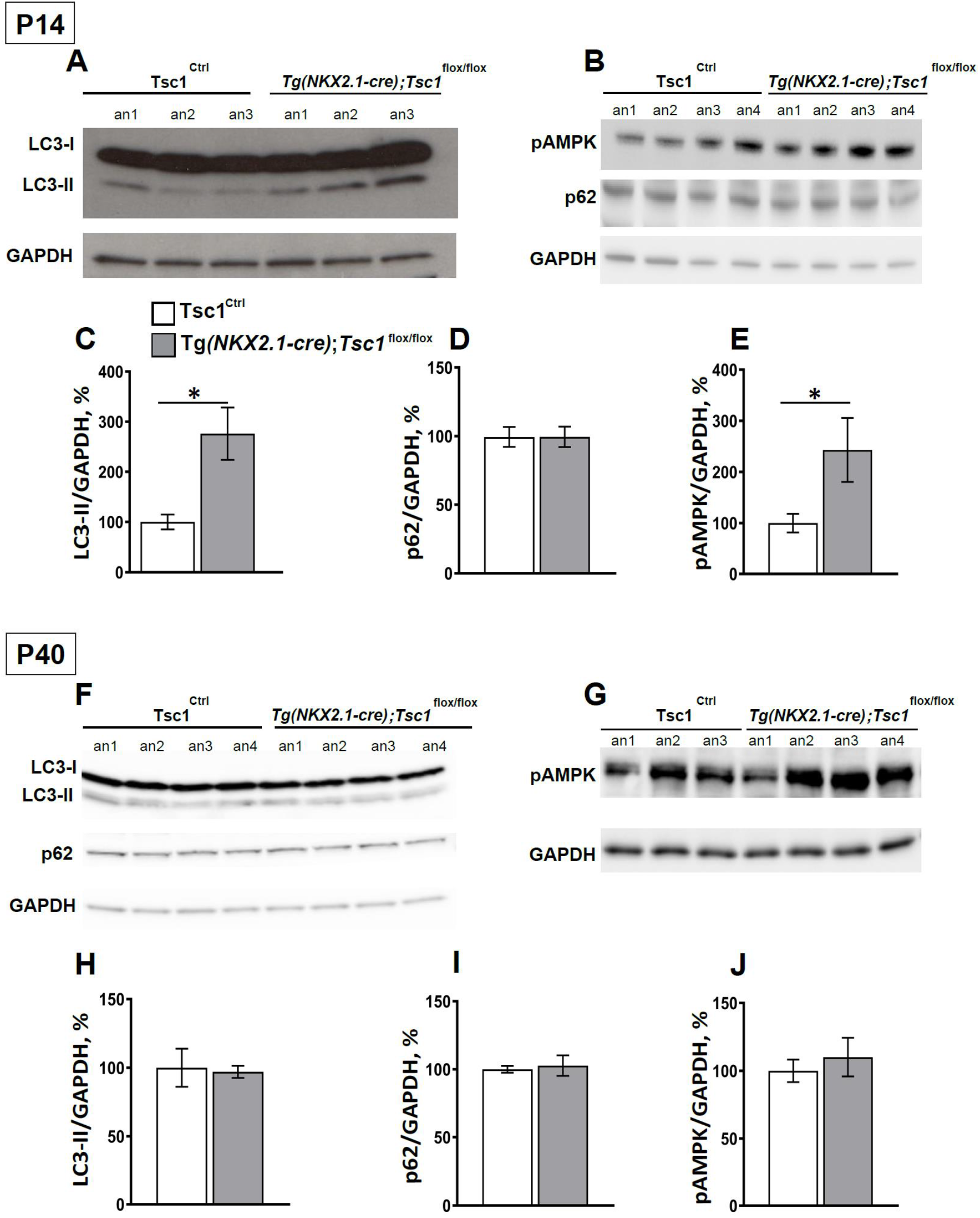
*Tsc1* deletion in GABAergic cells causes transient autophagy dysfunctions in adolescent mice. Western blot representative bands of LC3-I, LC3-II, p62 proteins (**A**) and pAMPK (**B**) and their quantification reveal that LC3-II and pAMPK expression are higher (**C**,**E**) in p14 Tg(*Nkx2.1-Cre*);*Tsc1^flox/flox^* **vs** *Tsc1*^Ctrl^ mice (Unpaired t-test: LC3-II *p=0.0314; pAMPK *p=0.0289), while p62 protein expression was unchanged (**D**) (Unpaired t-test: p=0,9937). **F, G,**Western blot representative bands of LC3-I, LC3-II, p62 (**F**) and pAMPK (**G**) in p40 mice and their quantification (**H, I, J**) show no differences between the two genotypes (Unpaired t-test: LC3-II p=0.8446; p62 p=0,7426; pAMPK p=0.6054). Number of mice at p14: LC3-II; *Tsc1*^Ctrl^ n=3, Tg(*Nkx2.1-Cre*);*Tsc1^flox/flox^* n= 3; p62 and pAMPK Tsc1^Ctrl^ n = 7, Tg(*Nkx2.1-Cre*);*Tsc1^flox/flox^* n= 5. Number of mice at p40: LC3-II and p62; Tsc1^Ctrl^ n = 4, Tg(*Nkx2.1-Cre*);*Tsc1^flox/flox^* n= 4; pAMPK; Tsc1^Ctrl^ n = 3, Tg(*Nkx2.1-Cre*);*Tsc1^flox/flox^* n= 4. Data represent mean ± SEM.

### Short-term administration of Rapamycin rescues long-term loss of PV cell innervation in Tg(*Nkx2.1-Cre*);*Tsc1^flox/+^* mice

So far, our data suggests that *Tsc1* haploinsufficiency in PV cells induces a premature formation of PV perisomatic synapses, which are however not stable and are subsequently lost. We can conceive two mechanisms to explain our observations: 1) Tsc1-mTORC1 signaling plays two distinct, opposing and age-dependent roles in PV cells, namely, during the first few postnatal weeks, mTORC1 activation promotes PV cell synapse formation, while later it may actively promote synapse pruning, or 2) mTORC1 hyper-activation during an early postnatal phase promotes PV cell synapse formation and can cause alterations (such as increased AMPK activation and autophagy) that are directly responsible for the synaptic loss occurring at later ages. To test which one of these two mechanisms is more likely to play a role in the loss of adult PV cell connectivity caused by *Tsc1* deletion, we biolistically transfected PV cells in EP26 cortical organotypic cultures prepared from *Tsc1^flox/flox^* mice with GFP (P_G67_-GFP/Cre) or GFP alone (control) and analyzed them at EP34 (Fig.6). We reasoned that if the first scenario was more likely, then late-onset Tsc1-deletion in PV cells should still cause loss of PV cell innervations. However, our data showed that the innervations formed by *Tsc1^−/−^* PV cells are indistinguishable from those formed by age-matched control PV cells for all analyzed parameters, suggesting that Tsc1-mTORC1 dysregulation before the third postnatal week is likely responsible for the subsequent loss of PV cell connectivity.

This observation raised the possibility that inhibiting mTORC1 hyperactivation during this critical time window might be sufficient to lead to long-term rescue of PV cell connectivity and, possibly, social behavior deficits. To directly test this hypothesis, we first used an *in vitro* approach by treating organotypic cultures from heterozygous and homozygous mutant mice with rapamycin (90ng/ml) from EP10 to EP18 and analyzing PV cell innervation at EP34. In culture from heterozygous mutant mice, rapamycin treatment reversed the decrease in perisomatic bouton density (Fig.S9E), terminal axonal branching (Fig.S9F) and percentage of target cell innervations (Fig.S9G) caused by *Tsc1* haploinsufficiency. We noted that rapamycin treatment also significantly reduced the percentage of target cells potentially contacted by PV cells in wild-type cultures (Fig.S9G).

In cultures from homozygous mutant mice, the same rapamycin treatment only partially reversed the decrease in perisomatic bouton density and terminal axonal branching, while it had not significant effect on the percentage of innervation formed by the mutant PV cells (Suppl Figure 10). It is possible that higher rapamycin doses might be required to completely rescue the PV cell innervation phenotype in PV cells from conditional homozygous mice.

Taken together, these data suggest that short-term rapamycin treatment during the early postnatal development can lead to persistent rescue of PV cell connectivity, particularly in case of haploinsufficiency.

Finally, to test whether short-term rapamycin treatment during early postnatal development can rescue long term effects of *Tsc1* haploinsufficiency *in vivo*, we treated *PV-Cre; Tsc1^flox/+^, PV-Cre;Tsc1^flox/flox^* and control littermates daily with either rapamycin (3 mg/kg; i.p.) or vehicle from P14 to P21 and analyzed PV cell perisomatic synaptic density and social behavior at P45 (Fig.8A). Interestingly, we found that rapamycin treatment restricted during this specific early developmental time window was sufficient to rescue the density of PV+/Geph+ puncta to wild-type levels in both hetero and homozygous mutant mice (Fig.8 B-I). In contrast, rapamycin treatment completely rescued social behavior deficits (both social approach and social novelty preference) in conditional heterozygous (*PV-Cre; Tsc1^flox/+^*) but not in conditional homozygous mice (*PV-Cre; Tsc1^flox/flox^*) (Fig.8L, M).

**Figure 7.**
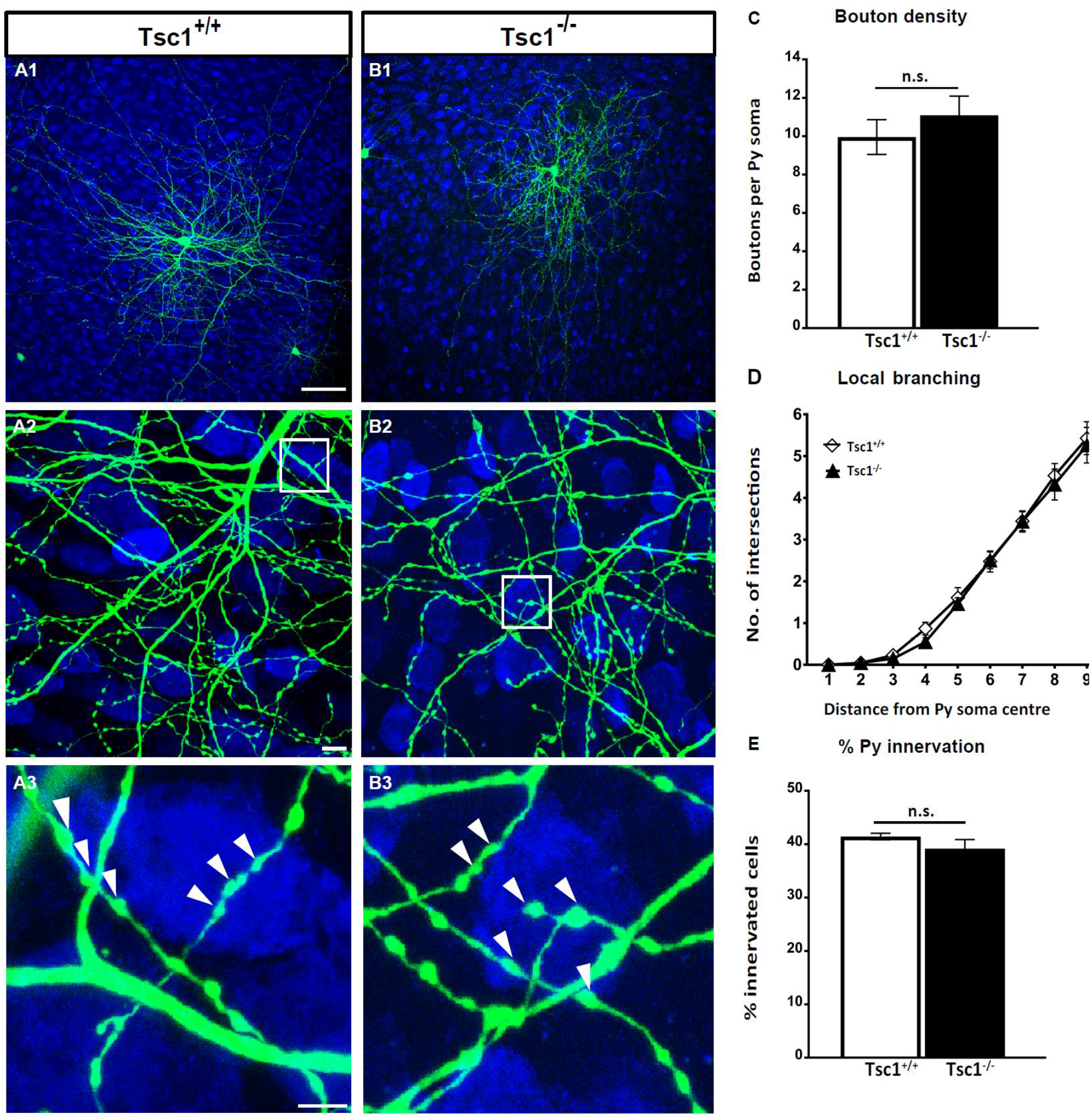
Late-onset *Tsc1* deletion in PV cells does not affect their innervation. **A1,** EP 34 *Tsc1*^+/+^ and **B1**, *Tsc1^−/−^* PV cells show similar axonal branching (**A2, B2**) and perisomatic boutons (**A3, B3,** arrowheads). **C,** Bouton Density (Welch’s t test, p= 0.4091), **(D)** local branching and (**E**) percentage of innervation (Welch’s t test, p=0.2448) are not significantly (n.s.) different between the two groups. Scale bars: ***A1-B1*** 50 μm; ***A2-B2*** 10 μm and ***A3-B3***, 5 μm. n=9 *Tsc1*^+/+^ PV cells, n=11 *Tsc1^−/−^* PV cells. Data in ***C-E*** represent mean ± SEM.

**Figure 8.**
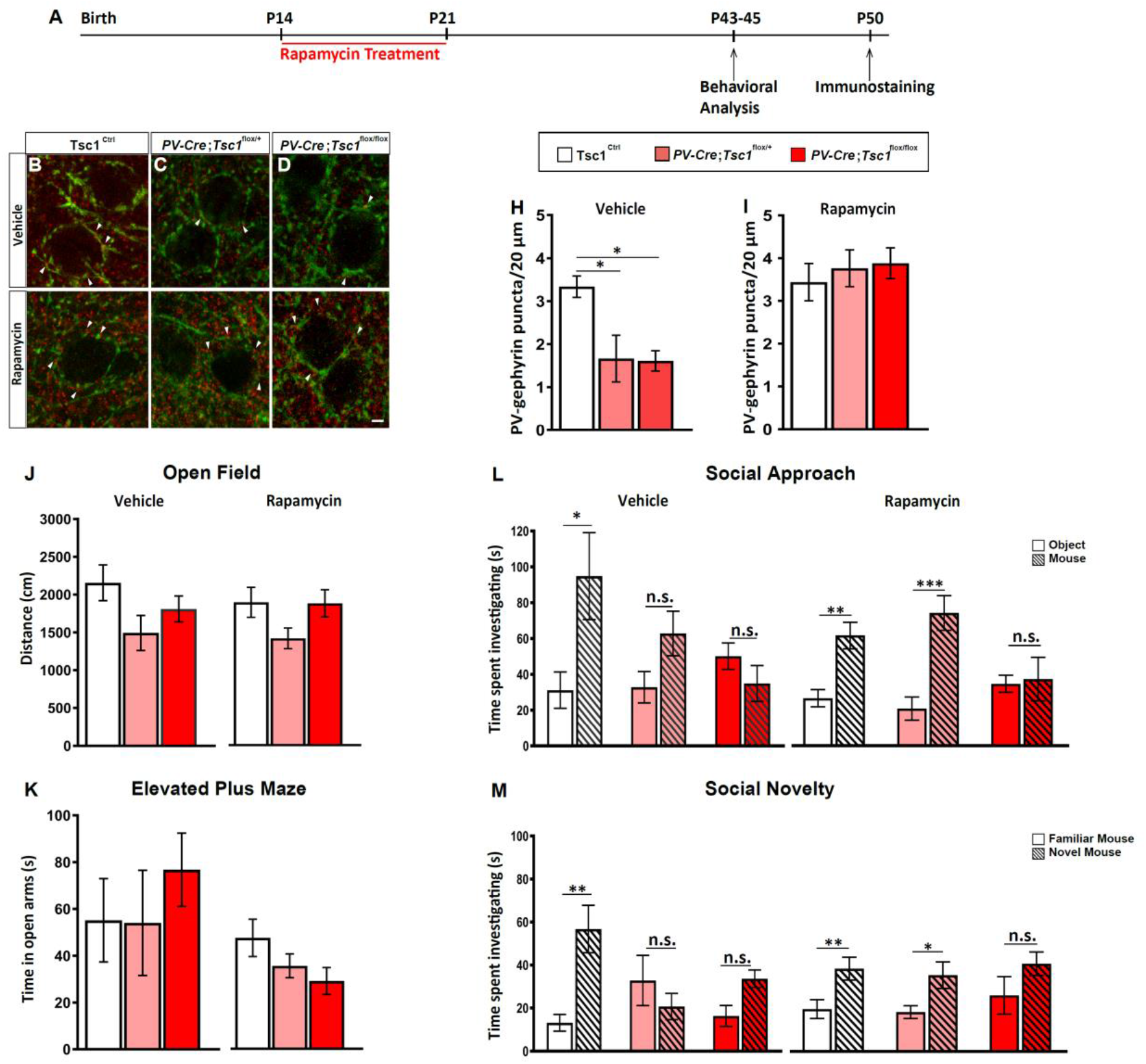
Short term Rapamycin treatment rescues loss of PV cell connectivity and social behavior deficits in adult heterozygous mutant mice. **A**, Schematic for treatment paradigm. **B-G**, Representative immunostained sections of somatosensory cortex labeled for PV (green) and gephyrin (red) in Vehicle (**B-D**) and Rapamycin (**E-G**) treated mice showing PV-Gephyrin colocalized boutons (arrowheads). **H**, PV-Gephyrin colocalized puncta in Vehicle (One-Way Anova, *p=0.036; Tukey’s multiple comparisons test: *Tsc1*^Ctrl^ **vs** *PV-Cre;Tsc1^flox/+^* *p=0.0401; *Tsc1*^Ctrl^ **vs** *PV-Cre;Tsc1^flox/flox^* *p=0.0281, *PV-Cre;Tsc^flox/+^* **vs** *PV-Cre;Tsc1^flox/flox^* *p=0.9942). Number of mice: *Tsc1*^Ctrl^, n=3; *PV-Cre;Tsc1^flox/+^*, n=4; *PV-Cre;Tsc1^flox/flox^* n=5 and (**I**) Rapamycin treated mice (One-Way Anova, p=0.7355; Tukey’s multiple comparisons test: *Tsc1*^Ctrl^ **vs** *PV-Cre;Tsc1^flox/+^* p=0.8421; *Tsc1*^Ctrl^ **vs** *PV-Cre;Tsc1^flox/flox^* p=0.7298, *PV-Cre; Tsc1^flox/+^* **vs** *PV-Cre;Tsc1^flox/flox^* p=0.9774). Number of mice: n=5 for all the genotypes. **J**, Vehicle (number of mice: *Tsc1*^Ctrl^, n=10; *PV-Cre;Tsc1^flox/+^* n=7; *PV-Cre;Tsc1^flox/flox^*, n=5) and Rapamycin (number of mice: *Tsc1*^Ctrl^, n=17; *PV-Cre;Tsc1^flox/+^*, n=10; *PV-Cre;Tsc1^flox/flox^*, n=10) treated mice travelled the same distance in Open Field and spent the same amount of time in the opens arms of the EPM (**K**). Number of mice; Vehicle: *Tsc1*^Ctrl^, n=16; *PV-Cre; Tsc1^flox/+^*, n=7; *PV-Cre;Tsc1^flox/flox^*, n=9; Rapamycin: *Tsc1*^Ctrl^, n=17; *PV-Cre;Tsc1^flox/+^*, n=11; *PV-Cre;Tsc1^flox/flox^*, n=11. **L,M,** Rapamycin treatment initiated at P14 rescues social approach (**L**, two-way ANOVA with Holm-Sidak’s post hoc analysis, *p<0.05, **p<0.001, ***p<0.0001) and social novelty deficits (**M**, two-way ANOVA with Holm-Sidak’s post hoc analysis, *p<0.05, **p<0.001, ***p<0.0001) in *PV-Cre; Tsc1^flox/+^* rapamycin treated mice but fails to rescue *PV-Cre;Tsc1^flox/flox^* mice. Number of mice; Vehicle: *Tsc1*^Ctrl^, n=7; *PV-Cre;Tsc1^flox/+^*, n=9; *PV-Cre;Tsc1^flox/flox^*, n=7; Rapamycin: *Tsc1*^Ctrl^, n=14; *PV-Cre;Tsc1^flox/+^*, n=10; *PV-Cre;Tsc1^flox/flox^*, n=8. Scale Bar: 5 μm. Data represent mean ± SEM.

Since PV is also expressed in Purkinje and granule cells in the cerebellum and Purkinje cellspecific deletion of *Tsc1* has been shown to cause social behavioral deficits (Tsai et al., 2012), we looked at cerebellum cyto-architecture, by immunolabeling cerebellar slices from vehicle- and rapamycin-treated mice with Calbindin, PV and NeuN (Suppl Figure 11). While we did not observe any obvious abnormalities in *PV-Cre;Tsc1^flox/+^* mice, the cerebellar cellular anatomy of *PV-Cre;Tsc1^flox/flox^* mice was significantly altered. In particular Purkinje cell numbers were greatly reduced and their dendritic arbors severely abnormal, resembling those observed during the first postnatal week. These abnormalities were only partially rescued by rapamycin treatment from P14-21. Thus, cerebellar impairments might contribute to the lack of rescue of social behavior deficits by early rapamycin treatment in the conditional homozygous mice (Tsai et al., 2018, Tsai et al., 2012).

In summary, taken together, the rescue data suggests that short-term rapamycin treatment during a critical postnatal window has long-lasting protective effects on GABAergic connectivity and social behavior in the context of *Tsc1* haploinsufficiency, which is typical of TSC patients.

## DISCUSSION

GABAergic circuits play a central role in the social behaviors affected in mTORC1-related neurodevelopmental disorders (Yizhar et al., 2011, Selimbeyoglu et al., 2017, Cao et al., 2018a). Here, we used single cell genetic manipulation approaches and genetic mouse models to investigate how dysregulation of mTORC1 signaling affects the development and maintenance of cortical PV GABAergic cells. We found that PV cell-specific haplosinsufficiency of *Tsc1*, a key negative regulator of mTORC1 signaling, leads to reduced PV cell connectivity, both at the input and output levels, since mutant PV cells formed less and smaller synapses and were contacted by less excitatory synapses. Mice either haploinsufficient or lacking *Tsc1* in PV cell exhibit altered social behavior. The fact that the conditional heterozygous mice exhibited comparable social behavior dysfunction as the conditional homozygous mice suggest that these findings are relevant for TSC, which is an autosomic dominant disorder.

Based on our data, we hypothesize that the loss of PV cell connectivity in adult mice is dependent on the premature formation of PV cell innervations during a critical, developmental period. First, single-PV cell *Tsc1* deletion with onset at EP10 caused a premature formation of PV cell innervation by EP18, followed by excessive synaptic pruning, while *Tsc1* deletion with onset at EP26, after PV cell innervations are stabilized, did not caused any changes in perisomatic innervations. In particular, the effects of *Tsc1* deletion in single, sparse PV cells in otherwise wild-type organotypic cultures suggest that *Tsc1* acts in a cell autonomous fashion to regulate PV cell innervation. Second, one-week rapamycin treatment from EP10-18 was sufficient to protect long-term PV cell innervation from excessive pruning. Third, consistent with the data *in vitro*, deletion or haploinsufficiency of *Tsc1 in vivo* leads to premature increases of PV cell perisomatic innervations in preadolescent mice followed by a significant loss in adults. On the other hand, mutant mice treated with rapamycin during a restricted, sensitive period (P14-22) did not show any loss of perisomatic PV cell putative synapses in adults compared to wild-type littermates. Fourth, conditional mutant *Tsc1* mice showed altered autophagy-associated processes at P14, when PV cells are at the peak of their maturation phase (Chattopadhyaya et al., 2004, Baho and Di Cristo, 2012, Baho et al., 2019, Okaty et al., 2009) but not at 6 postnatal weeks, when PV cell connectivity has already reached maturity.

Multiple, parallel cellular mechanisms most likely underlie the altered developmental time course of PV cell connectivity. *Tsc1* deletion-mediated mTORC1 hyperactivity may promote growth, via increased protein synthesis (Saxton and Sabatini, 2017a, Costa-Mattioli and Monteggia, 2013). In addition, mTORC1 activation has been shown to affect autophagy. Importantly, autophagy is mechanistically distinct in neurons compared to dividing cells. In fact, most studies on mTORC1 function, which used dividing cells, concluded that mTORC1 activation inhibits autophagy. On the other hand, Di Nardo and colleagues showed that *Tsc1/2*-deficient neurons displayed increased autophagic activity, which was dependent of AMPK activation (Di Nardo et al., 2014). Our results are consistent with these findings, since we observed increased AMPK phosphorylation and LC3-II levels in *Tsc1*-deficient GABAergic cells *in vivo*. Compromised autophagy, and accumulation of defective organelles, for example mitochondria, may be one of the downstream causes of PV cell axonal loss and synaptic pruning (Ebrahimi-Fakhari et al., 2016, Tang et al., 2014, Hui and Tanaka, 2019). Another critical factor controlling the maturation and maintenance of PV cell axonal arbour and synaptic boutons is their activity levels (Baho and Di Cristo, 2012, Stedehouder et al., 2018), which is modulated by their synaptic inputs. Adult PV cells in *PV-Cre;Tsc1^lox/+^* mice showed significantly reduced density of asymmetrical, presumably excitatory, synapses onto their dendrites, which may further contribute to reduced spiking activity and loss of PV cell innervations. Further studies are needed to dissect the relative contributions of these different mechanisms to the cellular phenotypes of mutant PV cells.

A recent study showed that Tsc1 deletion specifically in somatostatin (SST)-expressing GABAergic interneurons leads to altered firing properties of a percentage of cortical SST neurons in both conditional heterozygous and homozygous mice but to overall reduced synaptic output only in the conditional homozygous mutants (Malik, Nat Comm 2019), suggesting that the formation and refinement of PV cell synaptic connectivity is more sensitive to *Tsc1* haploinsufficiency than SST neurons even if they both originate from the medial ganglionic eminence. Another group generated *PV-Cre; Tsc1* conditional knockout mice, but in contrast to our findings, found no physiological phenotypes (Zhao, Yoshii Mol Brain 2019), thus concluding that most TSC phenotypes arise from excitatory pyramidal neurons. One possible explanation for this discrepancy is that in this study electrophysiological analysis was performed at P28-P30, which is after the phase of premature synapse formation but before synaptic loss might become detectable in *Tsc1* haploinsufficient PV cells. Consistent with this hypothesis, we did not observe significant differences in perisomatic innervations formed by mutant *Tsc1* PV cells in organotypic cultures at EP24 (Suppl. Fig.6).

Our data suggest the existence of a sensitive period, namely a time window during which therapy is effective for the treatment of a specific phenotype (LeBlanc and Fagiolini, 2011), for the treatment of social behavior impairments caused by *Tsc1* haploinsufficiency in PV cells. Strikingly, treatment limited to one week in pre-adolescent mice (P14-22) was sufficient to rescue both cortical PV cell innervation and social behavior deficits in adult *PV-Cre;Tsc1^lox/+^* mice, which may be clinically relevant since TSC patients carry germ line-derived heterozygous mutations. On the other hand, rapamycin treatment was sufficient to rescue cortical PV cell connectivity but not behavioural deficits in homozygous mutant mice, most likely due to the persistence of cerebellar defects. In fact, work from the Sahin’s group showed that *Tsc1*-deletion or haploinsufficiency in Purkinje cells was sufficient to cause autistic-like phenotypes in mice (Tsai et al., 2012). Interestingly, these cerebellar-dependent social behaviour phenotypes could be rescued by continuous rapamycin treatment initiated either at P7 (Tsai et al., 2012) or at 6 postnatal weeks (Tsai et al., 2018) in homozygous, Purkinje cell-specific mutant mice. It is possible that the sensitive period to ameliorate the deficits of cell survival and excitability caused by *Tsc1* deletion in Purkinje cells may be well into adolescence, later than that sufficient to rescue the deficits in cortical GABAergic PV interneurons. Alternatively, the chronic presence of rapamycin might be needed to inhibit the physiological changes in adult mutant Purkinje cells.

Multiple brain regions and neuronal circuits likely contribute to the different cognitive tasks, which underlie social behaviours. Cortical PV cell activity modulates sensory responses (Cardin et al., 2009, Cao et al., 2018a), which are required for the development of normal social interaction behaviours (Orefice et al., 2016). Postmortem analysis of brains from ASD patients as well as animal models for ASD (such as Mecp2 and Shank3 mutants) revealed abnormalities in PV cell circuits in multiple brain regions, including primary sensory cortices (Mierau et al., 2016, Chao et al., 2010, Patrizi et al., 2019, Orefice et al., 2019, Selimbeyoglu et al., 2017, Zikopoulos and Barbas, 2013, Nelson and Valakh, 2015, Vogt et al., 2018, Tomassy et al., 2014, Hashemi et al., 2017, Fukuda et al., 2005, Filice et al., 2016). Targeting PV cell circuit impairments might thus be a rational approach to ameliorate social interaction problems, however the developmental and cellular processes that lead to PV cell dysfunction are likely dependent of the underlying aetiology.

Our results suggest that *Tsc1* haploinsufficiency in PV cells leads to defects in adult connectivity that can be rescued by targeted treatment during a well-defined postnatal sensitive period. Multiple proteins in mTOR signalling pathway are either high confidence ASD-causative genes or underlie disorders with high ASD comorbidity (O’Roak et al., 2012), therefore highlighting this pathway as a possible etiological hub for the disorder. Whether a similar altered developmental trajectory of PV cell circuit maturation might be common to different mTORpathies remains to be explored. Interestingly, a recent study by Thion and collaborators reported that two different prenatal immune challenges lead to premature maturation of PV cell connectivity followed by reduced PV cell inhibitory drive in adult somatosensory cortex, similarly to what we observed (Thion et al., 2019). It will be interesting to investigate whether maternal immune activation affects PV circuit development by impinging on the Tsc/mTOR pathways.

Finally, while rapamycin, a mTORC1 blocker, was effective in rescuing PV cell connectivity and social behaviour deficits in *PV-Cre;Tsc1^lox/+^* mice, we cannot exclude that *Tsc1* deletion might have additional effects independent of mTORC1 signalling, since mice carrying hyperactive mTORC1 in Purkinje cells were recently reported to display different behavioural alterations compared to Purkinje-cell specific Tsc1-lacking mice (Sakai et al., 2019). A better understanding of the complexity of mTORC1 signalling network regulation, feedback and compensatory loops, may lead to the discovery of new molecular drug targets.

The mTOR inhibitor everolimus is approved by FDA for the treatment of subependymal giant cell astrocytomas, angiomyolipomas and complex partial seizures in TSC patients. Several recent studies have addressed whether everolimus treatment could have positive effects on cognition and autistic behaviour in TSC patients, but have so far produced controversial results (Hwang et al., 2016, Kilincaslan et al., 2017, Krueger et al., 2017, Mizuguchi et al., 2019, Ess and Franz, 2019, Overwater et al., 2019). One important point is that children younger than 4 years old were excluded from these studies. Our data suggests that an early age of onset of the treatment might be critical to improve specific cognitive and behavioural long term outcomes.

## ACKNOWLEDGEMENTS

We thank Drs. Elsa Rossignol (CHU Ste. Justine, Montreal, Canada) and Dr. Fabrice Ango (CNRS, Montpellier, France) for their insightful suggestions and Dr. Guy Doucet (Université de Montréal, Montreal, Canada) for providing reagents for Electron Microscopy. We would like to thank Antônia Samia Fernandes do Nascimento for her technical assistance, the Comité Institutionnel de Bonne Pratiques Animales en Recherche (CIBPAR), all the personnel of the animal facility of the Research Center of CHU Sainte-Justine (Université de Montreal) and the Plateforme Imagerie Microscopique for their instrumental technical support. This work was supported by the Canadian Institutes of Health Research (G.DC), Canada Foundation for Innovation (G.DC), Canada Research Chair Program (G.DC), and Natural Sciences and Engineering Research Council of Canada (NSERC). C.A.A. is supported by NSERC fellowship.

## MATERIAL AND METHODS

### Animals

*Tsc1* floxed mice with loxP sites flanking exons 17 & 18 of *Tsc1* gene (*Tsc1^flox/flox^*) were purchased from Jackson Laboratories (Cat# 005680). Two separate driver mouse lines expressing Cre recombinase, (1) *Tg(Nkx2.1-Cre)* (Xu et al., 2008), (Jackson Laboratories, Cat# 008661) and (2) *PV-Cre* (Jackson Laboratories, Cat# 008069) (Runyan et al., 2010) were crossed to the *Tsc1* floxed mice and the respective progenies were backcrossed to generate the heterozygous, homozygous and control genotypes within the same litter. To control for the pattern of expression of Cre, we introduced the RCE allele using Gt(ROSA)26Sortm1.1(CAG-EGFP)Fsh/J mice (Jackson laboratories). The RCE line carries a loxP-flanked STOP cassette upstream of eGFP sequence within the Rosa26 locus. Removal of the loxP-flanked STOP cassette by Cre-mediated recombination allows promoter-specific downstream eGFP expression (Sousa et al., 2009). All mice were housed under standard pathogen-free conditions in a 12h light/dark cycle with *ad libitum* access to sterilized laboratory chow diet. Animals were treated in accordance with Canadian Council for Animal Care and protocols were approved by the Animal Care Committee of CHU Ste-Justine Research Center.

### Mice Genotyping

DNA was extracted from mouse tails and genotyped to detect the presence of Cre alleles and Tsc1 conditional and wild-type alleles. Polymerase chain reaction (PCR) was performed using 3 separate primers; F4536 (5’-AGGAGGCCTCTTCTGCTACC-3’), R4830 (5’-CAGCTCCGACCATGA AGTG −3’) and R6548 (5’-TGGGTCCTGACCTATCTCCTA-3’) with band sizes of 295bp for the wild-type and 480bp for the floxed allele. 3 separate primers were also used for detecting Cre in the *Tg(Nkx2.1-Cre)* breeding; F1 (5’-AAGGCGGACTCGGTCCACTCCG-3’), F2 (5’-AAGGCGGACTCGG TCCACTCCG-3’) and R1 (5’-TCGGATC CGCCGCATAACCAG-3’) which generated 550bp and 220bp (mutant and wild-type) bands. Primers for detecting Cre in *PV*-Cre breeding were F1 (5’-CAGCCTCTGTTCCACATACACTCC-3’), F2 (5’-GCTCAGAGCCTCCATTCCCT-3’) and R1 (5’-TCACTCGAGAGTACCAAGCAGGCAGGA GATATC-3’) which generated 400bp and 526bp (mutant and wild-type) bands. To detect the presence of RCE alleles, 3 separate primers namely, RCE-Rosa1(5’-CCCAAAGTCGCTCTGAGTTGTTATC-3’), RCE-Rosa2(5’GAAGGAGCGGGAGAAATGGATATG-3,) and RCE-Cag3(5’-CCAGGCGGGC CATTTACCGTAAG-3’) were used which generated 350bp and 550bp bands.

### Slice culture and biolistic transfection

Slice culture preparation was done as described previously (Chattopadhyaya et al., 2004). Postnatal day 4 or 5 (P4 or P5) mouse pups were decapitated, and brains were rapidly removed and immersed in ice-cold culture medium (DMEM, 20% horse serum, 1 mM glutamine, 13 mM glucose, 1 mM CaCl_2_, 2 mM MgSO_4_, 0.5 μm/ml insulin, 30 mM HEPES, 5 mM NaHCO_3_, and 0.001% ascorbic acid). Coronal brain slices of the occipital cortex, 400 μm thick, were cut with a Chopper (Stoelting, Wood Dale, IL). Slices were then placed on transparent Millicell membrane inserts (Millipore, Bedford, MA), usually three to four slices/insert, in 30 mm Petri dishes containing 0.75 ml of culture medium. Finally, the slices were incubated in a humidified incubator at 34°C with a 5% CO_2_-enriched atmosphere and the medium was changed three times per week. All procedures were performed under sterile conditions. Constructs to be transfected were incorporated into “bullets” that were made using 1.6μm gold particles coated with a total of ~50 μg of the DNA(s) of interest. These bullets were used to biolistically transfect slices by Gene gun (Bio-Rad, Hercules, CA) at high pressure (180 Ψ). In order to delete *Tsc1* in single PV cells in an otherwise wild-type background, we transfected organotypic slices from *Tsc1^flox/flox^* mice either with P_G67_-GFP (Tsc1^+/+^, control PV cells) or P_G67_-Cre/ P_G67_-GFP (Tsc1^−/−^ PV cells). Organotypic cultures from Tg(*Nkx2.1-Cre^+/−^;Tsc1^flox/+^*), Tg(*Nkx2.1-Cre^+/−^;Tsc1^flox/flox^*) and Tsc1^Ctrl^ were transfected with P_G67_-GFP to visualize PV cells. For each experimental group, cortical slices were prepared from at least three mice. The majority of neurons labeled by using the P_G67_ promoter were PV-positive cells (Chattopadhyaya et al., 2004, Chattopadhyaya et al., 2007, Chattopadhyaya et al., 2013), while a minority (~10%) were pyramidal cells. Pyramidal cells were recognized by the complexity of their dendritic arbor, including an apical dendrite, and the presence of numerous dendritic spines. PV immunolabeling (see protocol below) was performed to confirm PV cell identity before imaging.

### Immunohistochemistry

Mice were perfused transcardially with saline followed by 4% Paraformaldehyde (PFA 4%) in phosphate buffer (PB 0.1M, pH 7.2). Brains were post-fixed with 4% PFA overnight and subsequently transferred to a 30% sucrose solution in sodium phosphate-buffer (PBS) for 48hrs. They were then frozen in molds filled with Tissue Tek using a 2-Methypentane bath cooled with a mixture of dry ice and ethanol (~70°C). optimal cutting temperature and coronal sections of 40 μm were obtained using a cryostat (Leica VT100). Organotypic cultures were fixed overnight at 4°C in 4% PFA in PB 0.1M, pH 7.2, then washed in PBS, incubated in 30% sucrose/PBS, and subjected to a freeze/thaw cycle at −20°C. Brain sections or organotypic cultures were blocked in 10% normal goat serum (NGS) and 1% Triton X-100 for 2 h at RT. Slices were then incubated for 48h at 4°C with the following primary antibodies: rabbit anti-phospho-S6 (1:1000, Cell Signaling, Cat# 5364), mouse anti-NeuN (1:400, Millipore, Cat# MAB377), chicken anti-NeuN (1:500, Millipore, Cat# ABN91), mouse anti-PV (1:1000, Millipore, Cat# 235), rabbit anti-PV (1:1000, Swant, Cat# PV27), guinea pig anti-PV (1:1000,Synaptic Systems, Cat# 195004), mouse anti-gephyrin (1:500, Synaptic Systems, Cat# 147021), mouse anti-Calbindin (1:1000, Abcam). It was followed by incubation with secondary antibodies for 2h at RT to visualize primary antibodies. The secondary antibodies used were Alexa-fluor conjugated 488, 555, 594, 633, and 647 (1:400, Life technologies; 1:1000, Cell Signaling Technology). After rinsing in PBS (three times), the slices were mounted in Vectashield mounting medium (Vector).

### Confocal Imaging and Quantitative analysis

All imaging was performed using Leica confocal microscopes (SPE, SP8 or SP8-STED). For PV cell innervation analysis, PV cells were first imaged using 10x (NA0.4) to record the overall cell morphology, and then multiple (2-3) stacks of their axonal fields were acquired using a glycerol immersion 63x (NA1.3) objective at 0.5 or 1 μm z-step in the first 150μm from the PV cell soma. The complexity of the PV axon branches around a pyramidal cell soma was reported as the average number of intersections, bouton density and percentage of innervated pyramidal cells. The number of intersections represented the intersections between a basket cell axon and the Sholl spheres (9 μm, increment of 1μm) from the center of the pyramidal cell soma. Bouton density around each basket cell represented the total number of GFP+ boutons in a radius of 9μm from the centre of the pyramidal cell soma. 12-24 pyramidal cells were analyzed for each basket neuron. To determine the percentage of pyramidal innervated by basket cells axon, we quantified the number of pyramidal cells soma that were contacted by the GFP+ axon and divided the later by the total number of somata in a confocal stack field.

For analysis *in vivo*, we imaged somatosensory cortex layers 2/3 and 5/6 using 20X (NA 0.75) and 63X oil (NA1.3) objectives. The 20X objective was used to acquire images for analyzing the percentage of PV/pS6 cellular colocalization, PV+/NeuN+ cell density, GFP+/GFP+PV+ cell density (specificity of PV recombination) and PV+/GFP+PV+ cell density (recombination rate). For the analysis of perisomatic innervation 63x glycerol objective was used to acquire images for quantifying PV and gephyrin puncta. At least three confocal stacks from 3 different brain sections were acquired in layers 2/3 and 5/6 of somatosensory cortex with z-step sizes of 0.5 (for synapse quantification) or 1μm (for cell density quantification). Cell soma size and cell density were quantified using Neurolucida (MBF Software). Fluorescence intensity of pS6 signal in PV cells was calculated using ImageJ or LAS X (Leica Application Suite X) software. In LAS X, 8 to 10 cells were chosen on various focal plane and encircled by using the polygon tool. This process generated the mean gray values of each cell. On the other hand, the mean gray values of four spots without any staining from the same focal plane were used as background, which were removed in order to normalize the data. For each animal, three sections were used in order to minimize the variability across the different groups. PV+, gephyrin+, and PV+/gephyrin+ puncta were counted around NeuN positive somata after selecting the confocal plane with the highest soma circumference using Neurolucida software. At least 6-10 NeuN positive somata were selected in each confocal stack. Investigators were blind to the genotypes during the analysis.

### Electron microscopy

The electron microscopy was carried out on 2 groups, *Tsc1*^Crl^ and *PV-Cre;Tsc1^flox/+^* mice, at P45. Mice were anesthetized and perfused with 0.1M PBS (0.9% Nacl in PB 0,2M; pH 7.4) followed by 2,5% glutaraldehyde + 2% PFA in 0.1M PB, pH 7.4. Following perfusion, the brains were further fixed for 2 hours at room temperature (RT) in the perfusion solution. Transverse 50-μm-thick sections of the brain were cut in cooled PBS with a vibratome (Leica, VT1000S). They were stored at −20°C in antifreeze solution (40% PB, 30% ethylene glycol, 30% glycerol) until used. Sections were immersed in 0.1% borohydride (in PBS) for 15 min at room temperature (RT), washed in PBS, and processed freely floating following a pre-embedding immunoperoxidase protocol previously described (Tremblay et al., 2007). Briefly, after rinsing in PBS, sections were preincubated (1-hour) at RT in a protein blocking solution (Expose Rabbit-Specific HRP/DAB detection IHC Kit, Abcam, Cambridge, UK, ab80437). Then, the sections were incubated for 48 hours at 4°C with rabbit anti-PV (1:1000, Swant, Cat# PV27) in PBS containing 1% NGS, followed by wash (three times in PBS) and incubation for 45 min at RT, in goat anti-rabbit horseradish peroxidase (HRP) conjugate (Abcam, Expose Kit, Cat# ab80437). After rinsing in PBS, immunoreactivity was visualized with hydrogen peroxide in the presence of di-aminobenzodine (DAB Chromogen, Abcam Expose Kit, Cat# ab80437). Thereafter, sections were rinsed in PB, postfixed flat in 1% osmium tetroxide for 1 hour and dehydrated in ascending concentrations of ethanol (50%, 70%, 90%, 100%, and finally in ethanol anhydrous). They were then treated with propylene oxide and then impregnated in resin overnight (Durcupan ACM; Sigma) at RT, mounted on aclar embedding film (EMS, Hatfield, PA) and cured at 55°C for 48 hours. Areas of interest from the somatosensory cortex (layers 5/6) were excised from the embedded sections and glued to the tip of prepolymerized resin blocks. Ultrathin (50-70 nm) sections were cut with an ultramicrotome (Reichart UltracutS, Leica, Wetzlar, Germany), collected on bare 150 square-mesh copper grids (Electron Microscopy Sciences, Hatfield, PA), stained with lead citrate, and examined at 80 KV with a Philips CM100 electron microscope, equipped with an 8 MB digital camera (AMT XR80).

To analyze the electron microscopy data, cellular profiles were identified according to well established criteria (Tremblay et al., 2007). All PV labeled structures were classified in different categories such as: dendritic shafts, axons and axon terminals. All the subcellular profiles that were difficult to identify were classified as “unknown”. To provide a better appraisal of the frequency of each type of cellular elements displaying immunolabelling, about eighty to hundred micrographs were randomly taken at 25000X in each animal, corresponding to a total surface of ~ 2000 μm^2^. Labeled profiles were counted in all micrograph. Results were expressed as number of immunopositive profiles per 100 μm^2^ of neuropil then normalized over results from the control mice. The area of neuropil and synapses lengths were measured using Neurolucida (MicroBrighField).

### Western Blot

Whole lysate proteins were extracted from the olfactory bulb where GABAergic cells are highly enriched. The olfactory bulbs of *Tsc1*^Ctrl^ and Tg(*Nkx2.1-Cre*);*Tsc1^flox/flox^* mice were dissected at P14 and P40 and snap frozen in liquid nitrogen. The tissue was then incubated in lysis buffer (150mM sodium chloride, 1% Triton x-100, 0.5 % sodium deoxycholate, 0.1% SDS, 50mM TrisHCl, pH 8, 2mM EDTA supplemented with a protease inhibitor cocktail III (Calbiochem)). The concentration of total protein was measured using the Bradford assay (BioRad). Proteins were separated on Novex Tris-Glycine 16% or NuPage Bis-Tris 4-12% protein gels (Invitrogen) in SDS running buffer and were transferred to PVDF membranes (BioRad). The following primary antibodies were used: anti-LC3B (1:1000, Novus), anti-p62 (1:500, Proteintech), anti-pAMPK (1:800, Cell Signaling), and anti-GAPDH (1:5000, ThermoFisher). Bands were quantified using Image J software. The intensity of LC3, p62 and pAMPK bands was normalized over the intensity of the GAPDH band.

### Rapamycin treatment

For *in vitro* experiments, organotypic cultures were prepared from Tg(*Nkx2.1-Cre*);*Tsc1^flox^* mice and were treated with Rapamycin from equivalent postnatal day 10 (EP10) to EP18. Rapamycin (90ng/ml, LC Laboratories, Woburn, MA, U.S.A.) was dissolved in the culture medium, which was changed every 48 hrs. For each mouse, half of the organotypic cultures were treated with rapamycin while the other half remained in regular culture medium, hence allowing us to have internal controls.

For *in vivo* treatment, rapamycin was administered daily (3 mg/kg; i.p.) to *PV-Cre;Tsc1^flox^* pups from P14 to P21. Rapamycin stock solution (20 mg/ml in 100% ethanol) was stored at – 20°C. Before injection, stock solution was diluted in 5% Tween 80 and 5% polyethylene glycol 400 to a final concentration of 1 mg/ml rapamycin in 4% ethanol (Buckmaster and Wen, 2011).

### Mouse behavior tests

Investigators were blind to genotype during both testing and analysis. Mice of both sexes were used in all experiments.

#### Open Field

A mouse was placed at the center of the open-field arena and the movement of the mouse was recorded by a video camera for 10 min. The recorded video file was later analyzed with the SMART video tracking system (v3.0, Harvard Apparatus). To measure exploratory behavior, total distance travelled during the 10 minutes period, and the time spent in the centre versus the periphery was calculated. The open field arena was cleaned with 70% ethanol and wiped with paper towels between each trial.

#### Elevated plus maze

The apparatus consists of two open arms without walls across from each other and perpendicular to two closed arms with walls joining at a central platform. A mouse was placed at the junction of the two open and closed arms. Time spent in the open versus closed arms was video recorded for 5 min. Recordings were scored to measure time spent in open arms, closed arms and center regions respectively.

#### 3 chamber social approach and social novelty tests

Mice (P45-60) were placed in the middle of the central chamber and allowed to explore all the chambers for 10 min for habituation. After habituation, a wire cage containing an unfamiliar conspecific of the same sex and age (Stranger 1) was placed inside one chamber, while an empty wire cage was placed in the second chamber. Mice were allowed to freely explore the three chambers of the apparatus for 10 min. Social approach was evaluated by quantifying the time spent by the test mice with the object or the mouse in each chamber during the 10 min session. At the end of 10min, a new unfamiliar mouse of the same sex and age (Stranger 2) was placed in the previously unoccupied wire cage and the test mouse observed for an additional 10 min to assess social novelty. Social novelty was evaluated by quantifying the time spent by the test mouse with either the familiar mouse (Stranger 1) or the newer mouse (Stranger 2) in each chamber during the third 10 min session. Strangers 1 and 2 originated from different home cages and had never been in physical contact with the test mice or with each other. Mice that stayed for the full 10 minutes session in only one chamber were excluded from the analysis.

### Statistical analysis

Differences between 2 experimental groups was assessed using t-test for normally distributed data and t-test with Welch’s correction for not normally distributed data. Differences between 3 or more experimental groups were assessed with one-way ANOVA and *post hoc* comparison. For non-normally distributed data, nonparametric Kruskal–Wallis one-way ANOVA test was used. In experiments involving Rapamycin treatment and social behavior, two-way ANOVA with *post hoc* analysis was used. Cumulative distributions were analyzed using the Kolmogorov-Smirnov test. All bar graphs represent mean ± SEM. All the statistical analyses were performed using Prism 7.0 (GraphPad Software).

