## Supplemental Material for "Inhibition of mTOR during a postnatal critical sensitive window rescues deficits in GABAergic PV cell connectivity and social behavior caused by loss of *TSC1*"

*Corresponding author :

Graziella Di Cristo,

E-mail: graziella.dicristo@recherche-ste- justine.qc.ca.

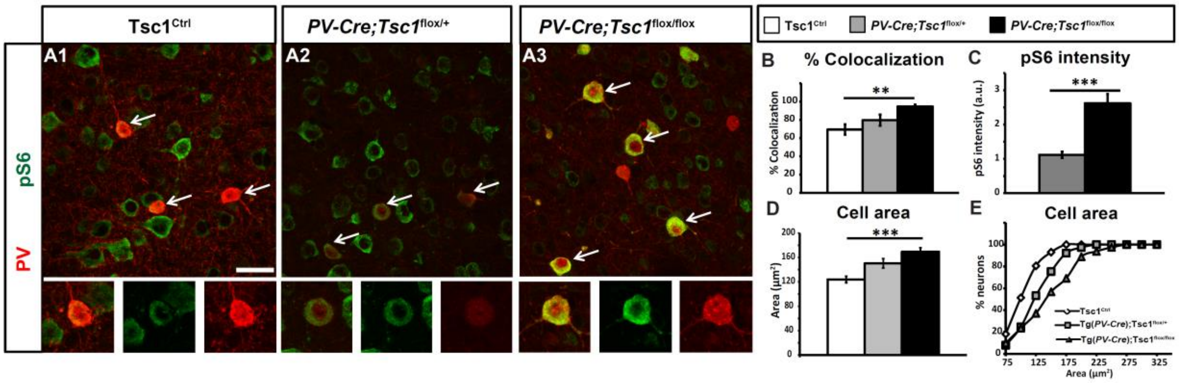


**Supplemental Figure 1. Cortical PV cells show increased mTOR activity and somatic hypertrophy in *PV-Cre;Tsc1*$^{flox/flox}$ mice.** **A**, Coronal sections of somatosensory cortex immunostained for PV (red) and pS6 (green) in *Tsc1*$^{Ctrl}$ (**A1**), *PV-Cre*;*Tsc1*$^{flox/+}$ (**A2**) and *PV-Cre*;*Tsc1*$^{flox/flox}$ mice (**A3**) at P45. Lower panels show individual PV cells. **B**, *PV-Cre*;*Tsc1*$^{flox/flox}$ mice show increased percentage of colocalization of pS6 in PV cells (one-way ANOVA, **p=0.0035; Holm-Sidak post hoc analysis: *Tsc1*$^{Ctrl}$ **vs** *PV-Cre*;*Tsc1*$^{flox/+}$ p=0.9097; *Tsc1*$^{Ctrl}$ **vs** *PV-Cre*;*Tsc1*$^{flox/flox}$ **p=0.0054). Number of mice: *Tsc1*$^{Ctrl}$ n = 4, *PV-Cre*;*Tsc1*$^{flox/+}$ n= 5, *PV-Cre*;*Tsc1*$^{flox/flox}$ n= 4. **C**, Quantification of pS6 expression intensity in PV cells normalized to wild-type controls show two-fold increase in *PV-Cre; Tsc1*$^{flox/flox}$ mice (Welch's t-test **p=0.0093). Number of mice: *PV-Cre;Tsc1*$^{flox/+}$ n= 7, *PV-Cre*;*Tsc1*$^{flox/flox}$ n= 5. **D, E,** PV cells show somatic hypertrophy in homozygous mutant mice. **D** (one-way ANOVA, ***p=0.0001; Dunnett post hoc analysis: *Tsc1*$^{Ctrl}$ **vs** *PV-Cre*;*Tsc1*$^{flox/+}$ p=0.6953; *Tsc1*$^{Ctrl}$ vs *PV-Cre*; *Tsc1*$^{flox/flox}$ ***p=0.0001). Number of mice: *Tsc1*$^{Ctrl}$ n = 6, *PV-Cre*; *Tsc1*$^{flox/+}$ n= 5, *PV-Cre*; *Tsc1*$^{flox/flox}$ n= 5. **E**, Cumulative distribution (K-S test, *p<0.001) at P45. Scale bar, 20 µm. Data represent mean ± SEM.

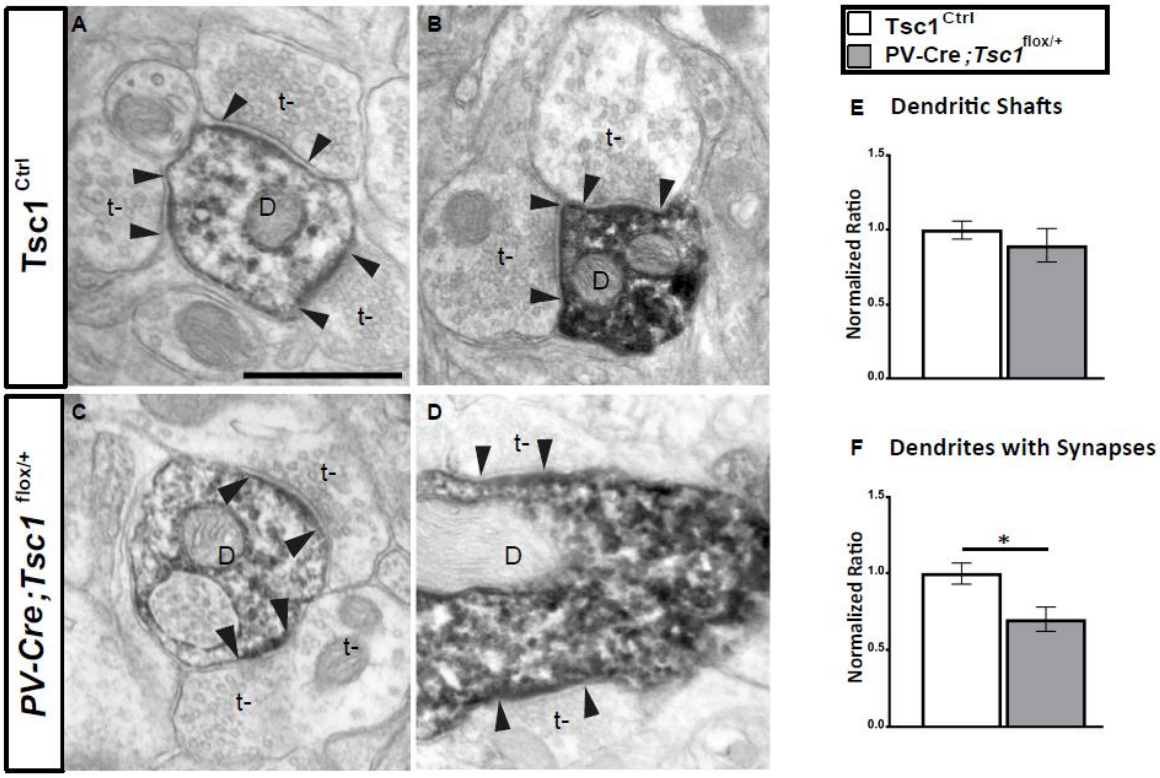


**Supplemental Figure 2. Tsc1 haploinsufficient PV cells show reduced density of asymmetric synapses onto their dendrites. A-D,** PV-immunolabeled dendritic branches in somatosensory cortex of *Tsc1*$^{\mathrm{Ctrl}}$ (**A**, **B**) and *PV-Cre;Tsc1*$^{flox/+}$ (**C**, **D**) at P60 showing multiple asymmetric synapses (arrowheads) between unlabeled axon terminal (t-) and a labeled dendritic shaft (indicated with a D in the microphotographs). **E**, **Overall** PV+ dendritic branch density is no different between the two genotypes (Welch's t-test, p=0.4665). **F**, Quantification of PV+ dendrites bearing synapses shows a significant decrease in *PV-Cre;Tsc1*$^{flox/+}$ mice compared to control littermates (Welch's t-test, *p=0.0422). Arrowheads indicate presence of PSDs (post synaptic density) in the dendrites. Number of mice: *Tsc1*$^{Ctrl}$, n=4; *PV-Cre*;*Tsc1*$^{flox/+}$, n=3. Scale bar, 500 nm.

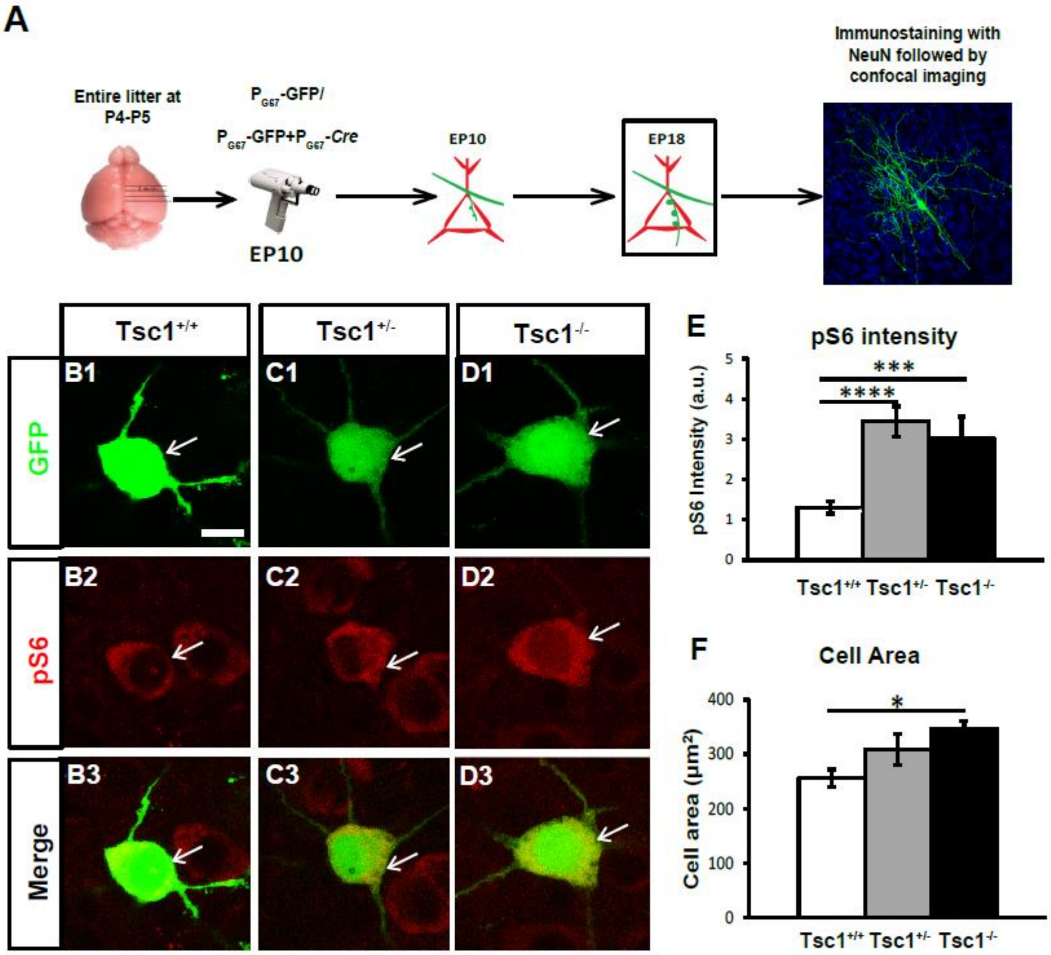


**Supplemental Figure 3. *Tsc1* knockout in single PV cells leads to increase in mTOR activity and somatic hypertrophy. A,** Schematics of experimental procedure. **B-D,** PV cells from cortical organotypic cultures transfected with $P_{G67}$ (*Tsc1*$^{+/+}$ control cells) or $P_{G67}$-Cre (*Tsc1*$^{+/-}$ and *Tsc1*$^{-/-}$) immunostained for pS6 (red) at EP18. **E**, Somatic pS6 intensity is increased in both *Tsc1*$^{+/-}$ (**C2**) and *Tsc1*$^{-/-}$ PV cells (**D2**) compared to *Tsc1*$^{+/+}$ PV cells (one-way ANOVA, ****p<0.0001; Holm-Sidak post hoc analysis: *Tsc1*$^{+/+}$ **vs** *Tsc1*$^{+/-}$ ****p<0.0001; *Tsc1*$^{+/+}$ **vs** *Tsc1*$^{-/-}$ ***p=0.0003). Number of cells: *Tsc1*$^{+/+}$ n = 17, *Tsc1*$^{+/-}$ n=7, *Tsc1*$^{-/-}$ n=7. ***F***, *Tsc1*$^{-/-}$ cells have increased soma area (one-way ANOVA, *p=0.0213; Holm-Sidak post hoc analysis: *Tsc1*$^{+/+}$ **vs** *Tsc1*$^{+/-}$ p=0.0876; *Tsc1*$^{+/+}$ **vs** *Tsc1*$^{-/-}$ *p=0.0185). Number of mice: *Tsc1*$^{+/+}$ n=20, *Tsc1*$^{+/-}$ n=9, *Tsc1*$^{-/-}$ n=7. Scale bar, 10µm. Data represents mean ± SEM.

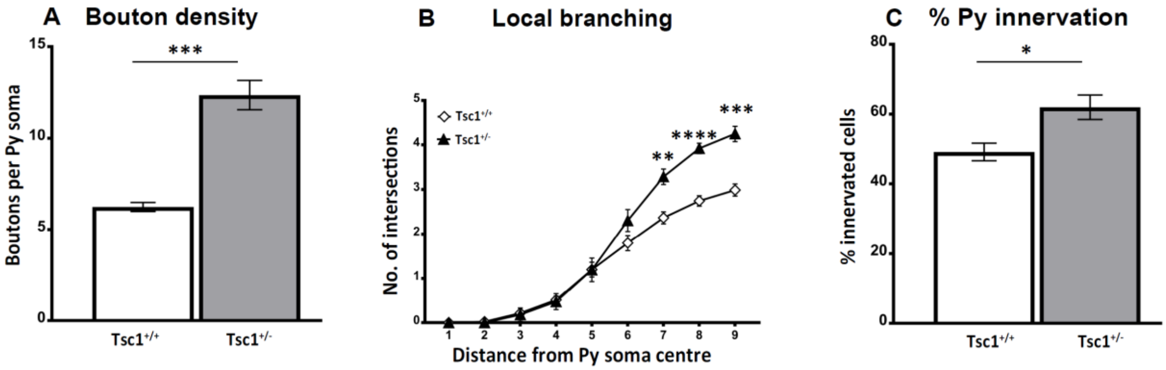


**Supplemental Figure 4. *Tsc1* haploinsufficiency in single PV cells causes a premature increase in axon terminal branching and bouton density. A**, PV cells lacking one allele of *Tsc1* show increase in bouton density (Welch's t-test, ***p=0.0004), number of PV cells: n=16 for *Tsc1*$^{+/+}$, n=6 for *Tsc1*$^{+/-}$ and local branching (**B**) (Welch's t-test, **p=0.0014 (radius7), ****p<0.0001 (radius8), ***p=0.0001 (radius9)), number of PV cells: n=14 for *Tsc1*$^{+/+}$, n=6 for *Tsc1*$^{+/-}$. **C**, Percentage of innervated cells (Welch's t-test, *p=0.0114). Number of PV cells: n=14 for *Tsc1*$^{+/+}$, n=7 for *Tsc1*$^{+/-}$. Data represent mean ± SEM.

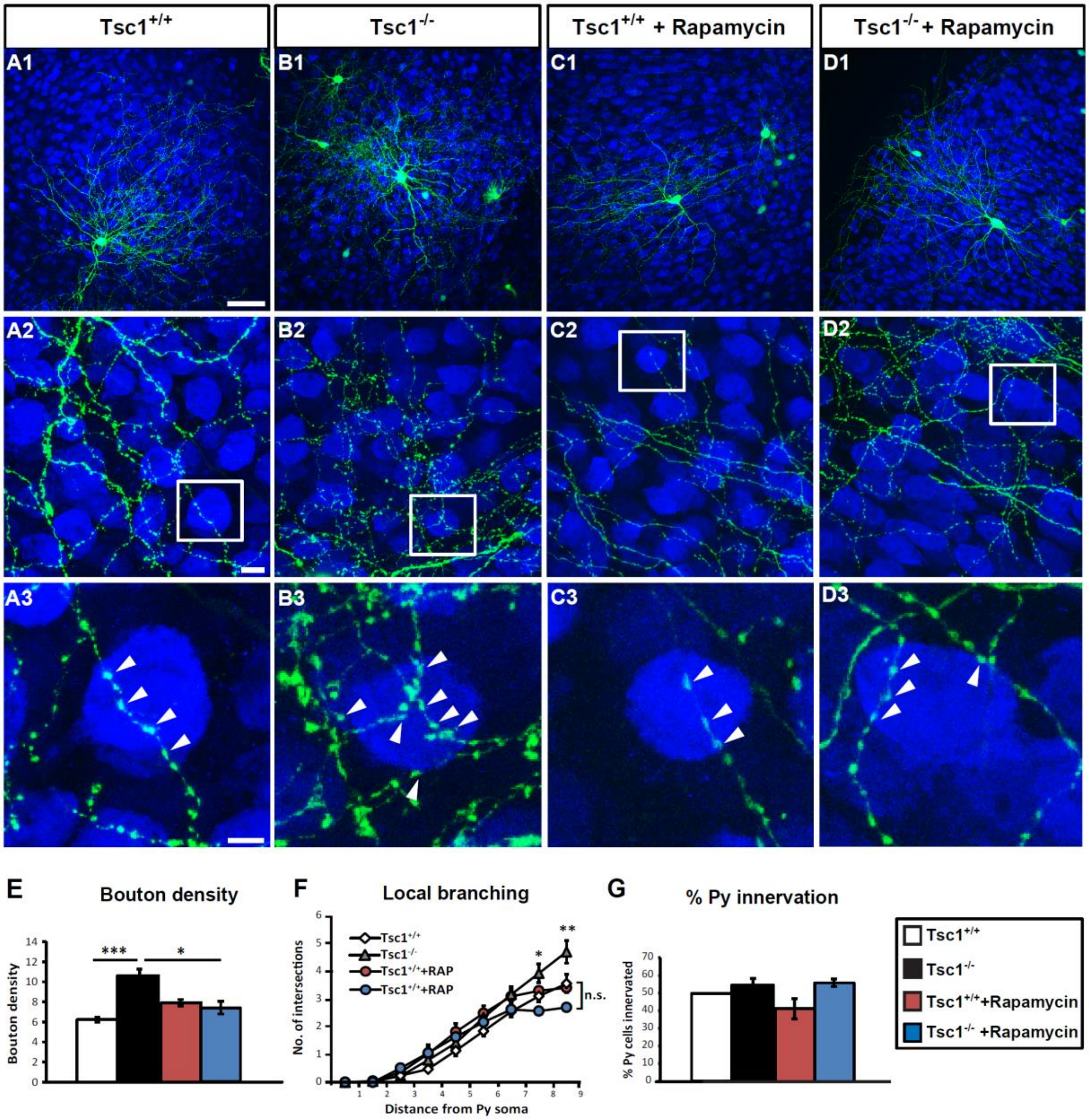


**Supplemental Figure 5. Premature increase in perisomatic innervation by *Tsc1*$^{-/-}$ PV cells is mTORC1 dependent. A, B**, *Tsc1*$^{-/-}$ PV cell (green) shows more complex terminal axonal branching (**A2, B2)** and increased bouton density at EP18 (**A3, B3,** arrowheads) compared to control, age-matched PV cells. **C, D:** Rapamycin treatment from EP12-18 does not affect bouton density and local branching of *Tsc1*$^{+/+}$ PV cells (**C**), while it normalizes perisomatic innervations formed by *Tsc1*$^{-/-}$ PV cells (**D)**. *Tsc1*$^{-/-}$ PV cells show increased bouton density (**E**) (two-way ANOVA with Bonferroni post hoc analysis, *p<0.05) and local branching (**F**) (two-way ANOVA with Bonferroni post hoc analysis, *p<0.05) compared to the other groups. **G**, Percentage of innervation. PV cells: n=9 *Tsc1*$^{+/+}$ PV cells, n=9 *Tsc1*$^{-/-}$ PV cells, n=4 *Tsc1*$^{+/+}$ + Rapamycin PV cells, n=4 *Tsc1*$^{-/-}$ + Rapamycin PV cells, Scale bars: ***A1-D1***, 100 µm; ***A2-D2*** and ***A3-D3***, 5 µm.

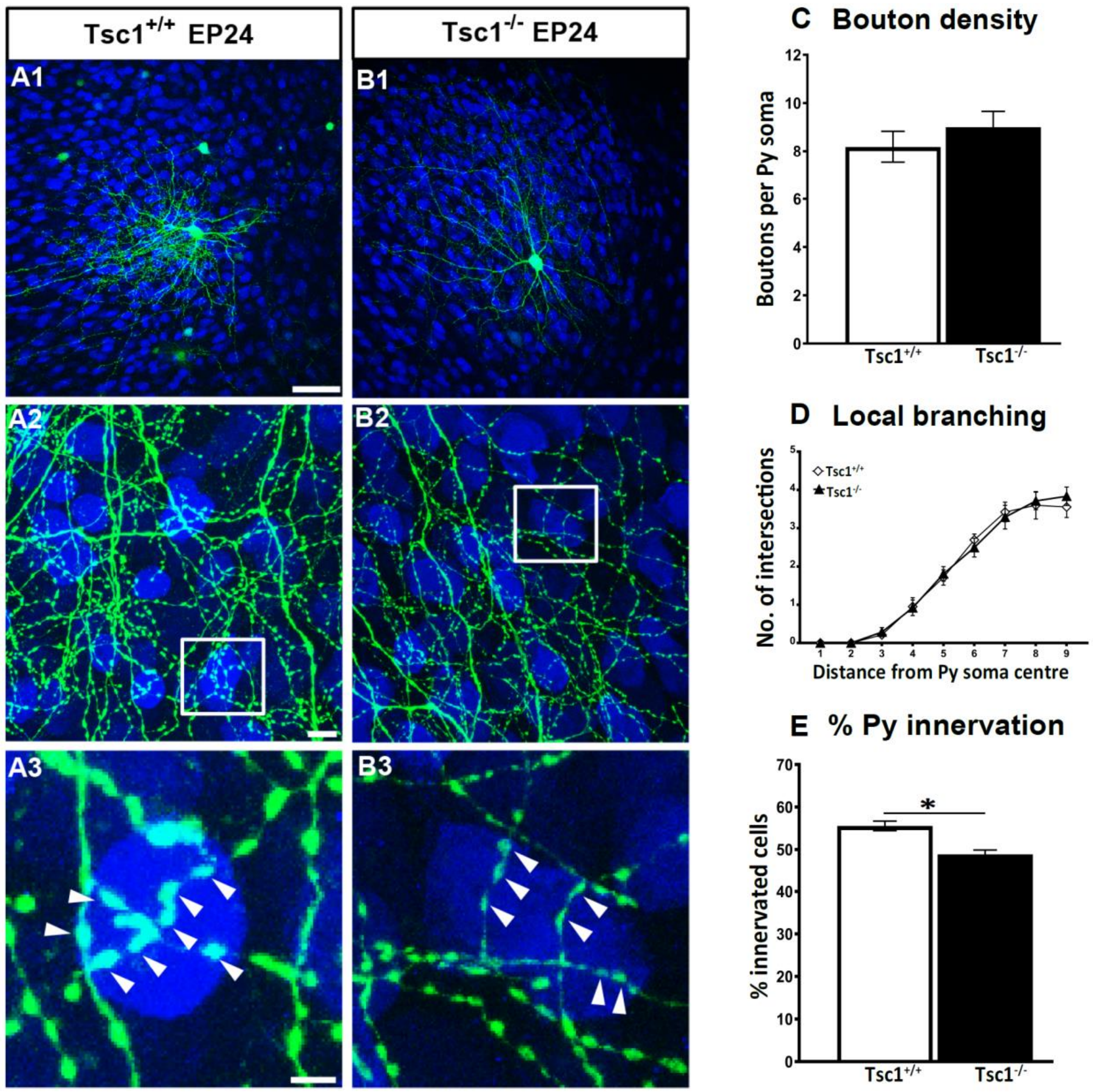


**Supplemental Figure 6. PV *Tsc1*$^{-/-}$ cell innervations at EP24 are morphologically indistinguishable from age-matched controls. A1**, *Tsc1*$^{+/+}$ and **B1**, *Tsc1*$^{-/-}$ PV cells show similar axonal branching (**A2, B2**) and perisomatic bouton density (**A3, B3,** arrowheads). **C**, Bouton density (Welch's t test, p=0.3920). **D,** local branching is not significantly different between the two groups. **H**, percentage of innervation is significantly reduced in the *Tsc1*$^{-/-}$ PV cells (Welch's t test ***p=0.0009). Number of PV cells: n=9 Ctrl *Tsc1*$^{+/+}$, n=6 *Tsc*$^{-/-}$ PV neurons. Scale bars: ***A1-B1***, 50 µm; ***A2-B2*** 10 µm, ***A3-B3***, 5 µm. Data represents mean ± SEM.

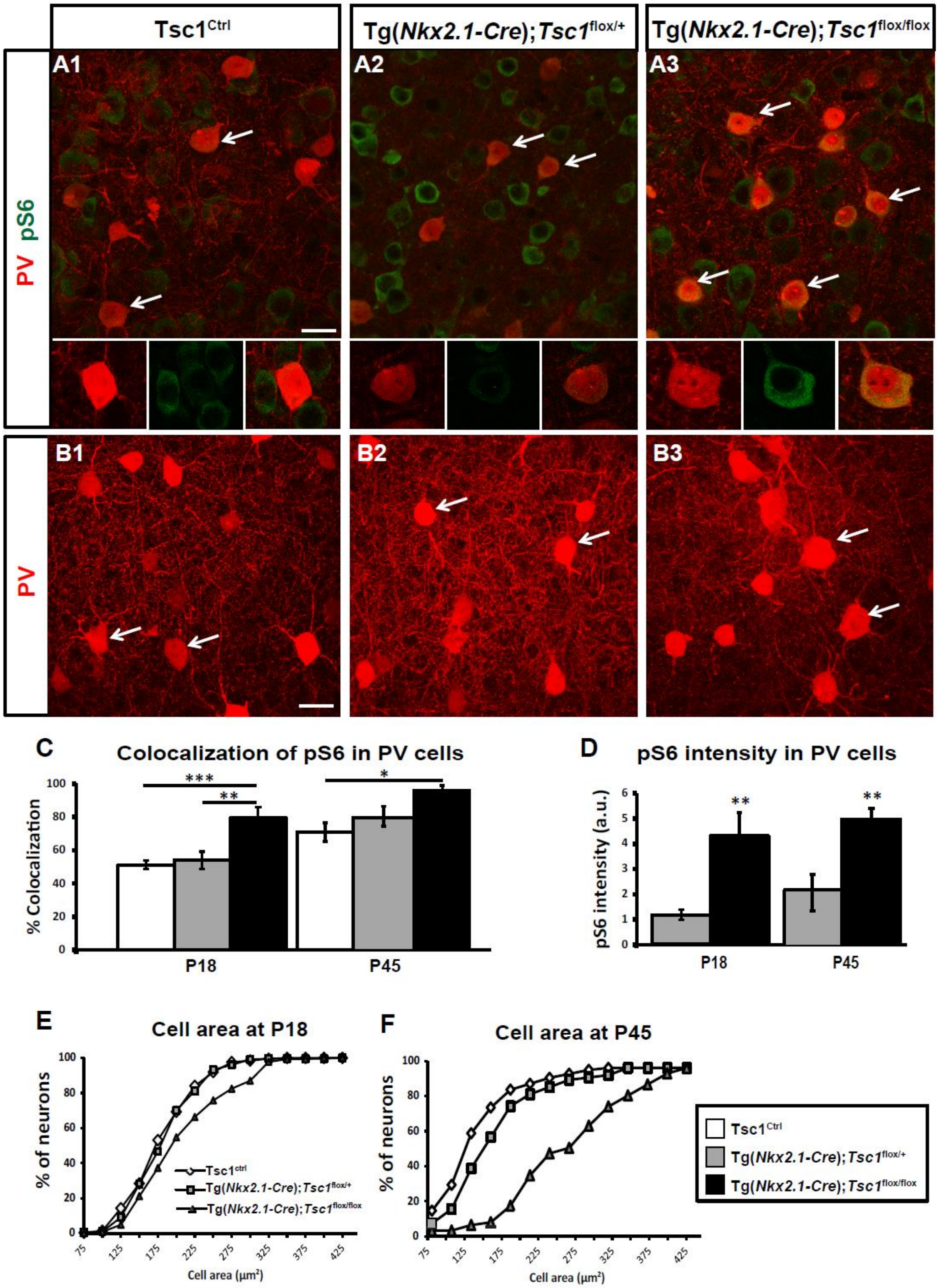


**Supplemental Figure 7. Cortical PV cells from Tg(*Nkx2.1-Cre*);*Tsc1*$^{flox/flox}$ mice show increased mTOR activity and somatic hypertrophy**. **A**, Coronal sections of somatosensory cortex immunostained for PV (red) and pS6 (green) (**A**) or PV only (**B**) in *Tsc1*$^{\mathrm{Ctrl}}$ mice (**A1, B1**), Tg(*Nkx2.1-Cre*);*Tsc1*$^{flox/+}$ mice (**A2, B2**) and Tg(*Nkx2.1-Cre*);*Tsc1*$^{flox/flox}$ mice (**A3, B3**) at P18. Lower panels show higher magnification of individual PV cells. **C**, In Tg(*Nkx2.1-Cre*); *Tsc1*$^{flox/flox}$ mice, more PV cells co-localize with pS6 as compared to Tg(*Nkx2.1-Cre*);*Tsc1*$^{flox/+}$ and wild-type mice at P18 (one-way ANOVA, ***p=0.0005; Tukey's multiple comparison test: *Tsc1*$^{\mathrm{Ctrl}}$ vs Tg*(Nkx2.1-Cre); Tsc1*$^{flox/+}$ p=0.9154; *Tsc1*$^{\mathrm{Ctrl}}$ **vs** Tg(*Nkx2.1-Cre); Tsc1*$^{flox/flox}$ ***p=0.0005; Tg*(Nkx2.1-Cre); Tsc1*$^{flox/+}$ **vs** *Tg(Nkx2.1-Cre); Tsc1*$^{flox/flox}$ **p=0.0072) and at P45 (one-way ANOVA, *p=0.0254; Tukey's multiple comparison test: *Tsc1*$^{Ctrl}$ vs *Tg(Nkx2.1-Cre); Tsc1*$^{flox/+}$ p=0.3805; *Tsc1*$^{\mathrm{Ctrl}}$ vs Tg*(Nkx2.1-Cre); Tsc1*$^{flox/flox}$ *p=0.0202; *Tg(Nkx2.1-Cre); Tsc1*$^{flox/+}$ *Tsc1*$^{\mathrm{Ctrl}}$ **vs** Tg*(Nkx2.1-Cre); Tsc1*$^{flox/flox}$ ***p=0.1789). Number of mice at P18 *Tsc1*$^{\mathrm{Ctrl}}$ n=11, Tg(*Nkx2.1-Cre); Tsc1*$^{flox/+}$ n=5, *Tg(Nkx2.1-Cre); Tsc1*$^{flox/flox}$ n=7; number of mice at P45, *Tsc1*$^{\mathrm{Ctrl}}$ n=5, Tg*(Nkx2.1-Cre); Tsc1*$^{flox/+}$ n=5, *Tg(Nkx2.1-Cre); Tsc1*$^{flox/flox}$ n=4. **D,** pS6 expression intensity in PV cells normalized to wild-type controls at P18 (t-test p<0.05; number of mice at P18, *Tg(Nkx2.1-Cre);Tsc1*$^{flox/+}$ n=6, *Tg(Nkx2.1-Cre); Tsc1*$^{flox/flox}$ n=6) and at P45 (t-test p<0.05; number of mice at P45, *Tg(Nkx2.1-Cre); Tsc1*$^{flox/+}$ n=6, Tg(*Nkx2.1-Cre*); *Tsc1*$^{flox/flox}$ n= 7. **E, F,** Quantification of PV cell area shows somatic hypertrophy in Tg(*Nkx2.1-Cre*); *Tsc1*$^{flox/flox}$ mice at both P18 and P45 (P18: K-S test, *p<0.01; P45: K-S test, *p<0.001), and in Tg(*Nkx2.1-Cre*); *Tsc1*$^{flox/+}$ mice at P45 (P18: K-S test, *p<0.05), n=11 *Tsc1*$^{\mathrm{Ctrl}}$ mice, n=5 Tg(*Nkx2.1-Cre*); *Tsc1*$^{flox/+}$ mice, n=7 Tg(*Nkx2.1-Cre*); *Tsc1*$^{flox/flox}$ mice at P18, n=6 mice for all genotypes at P45. Scale bar, 20 µm.

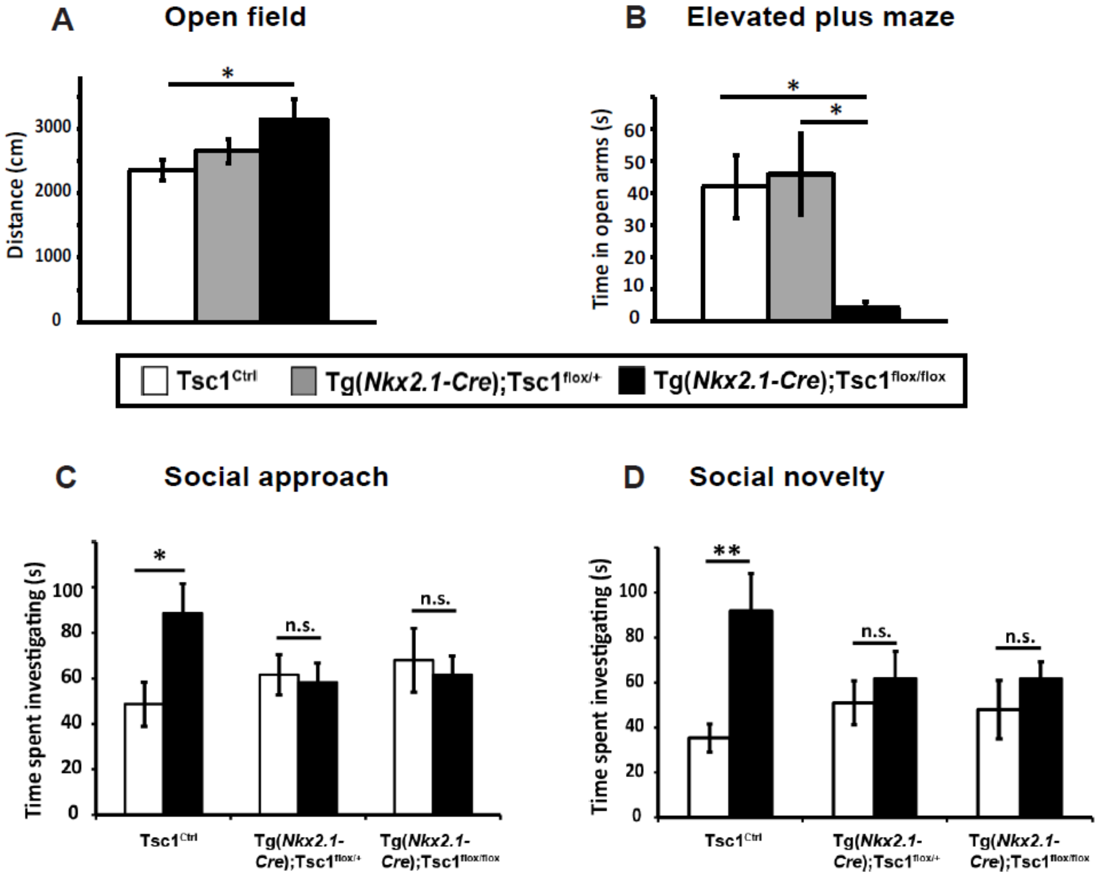


**Supplemental Figure 8. Tsc1 knockout in MGE derived neurons leads to social behavioral deficits in young adult mice. A**, Open field test: Quantification of distance travelled during exploratory activity in an open field arena at P33 shows increased exploratory drive in Tg*(Nkx2.1-Cre);Tsc1*$^{flox/flox}$ mice (one-way ANOVA, *p<0.0407; Holm-Sidak post-hoc analysis: *Tsc1*$^{\mathrm{Ctrl}}$ **vs** Tg*(Nkx2.1-Cre);Tsc1*$^{flox/+}$ p=0.3246; *Tsc1*$^{\mathrm{Ctrl}}$ vs Tg*(Nkx2.1-Cre); Tsc1*$^{flox/flox}$ *p=0.0241). Number of mice: *Tsc1*$^{\mathrm{Ctrl}}$ n=14, Tg*(Nkx2.1-Cre);Tsc1*$^{flox/+}$ n=10, Tg*(Nkx2.1-Cre); Tsc1*$^{flox/flox}$ n=10. **B**, Elevated plus maze: Quantification of time spent in the open arms of elevated plus maze arena at P35 shows increased anxiety like behaviour in Tg*(Nkx2.1-Cre); Tsc1*$^{flox/flox}$ mice (one-way ANOVA, *p=0.0137; Tukey's multiple comparisons test: *Tsc1*$^{\mathrm{Ctrl}}$ **vs** Tg*(Nkx2.1-Cre);Tsc1*$^{flx/+}$ p=0.7755; *Tsc1*$^{\mathrm{Ctrl}}$ **vs** Tg*(Nkx2.1-Cre); Tsc1*$^{flox/flox}$ *p=0.0230). Number of mice: *Tsc1*$^{\mathrm{Ctrl}}$ n=14, Tg*(Nkx2.1-Cre);Tsc1*$^{flox/+}$ n=12, Tg*(Nkx2.1-Cre); Tsc1*$^{flox/flox}$ n=10. **C**, In the 3 chambers, both Tg*(Nkx2.1-Cre);Tsc1*$^{flox/+}$ and Tg*(Nkx2.1-Cre); Tsc1*$^{flox/flox}$ mice do not show preference for the mouse vs the object (two-way ANOVA with Bonferroni's post hoc analysis, *p<0.05). Number of mice: *Tsc1*$^{\mathrm{Ctrl}}$ n=14, Tg*(Nkx2.1-Cre);Tsc1*$^{flox/+}$ n=10, Tg*(Nkx2.1-Cre); Tsc1*$^{flox/flox}$ n=12. ***D***, Unlike *Tsc1*$^{\mathrm{Ctrl}}$ mice, both Tg*(Nkx2.1-Cre);Tsc1*$^{flox/+}$ and Tg*(Nkx2.1-Cre); Tsc1*$^{flox/flox}$ mice failed to show preference for social novelty (two-way ANOVA with Bonferroni's post hoc analysis, **p<0.001). Number of mice: *Tsc1*$^{\mathrm{Ctrl}}$ n=14, Tg*(Nkx2.1-Cre);Tsc1*$^{flox/+}$ n=14, Tg*(Nkx2.1-Cre); Tsc1*$^{flox/flox}$ n=12. Data represents mean ± SEM.

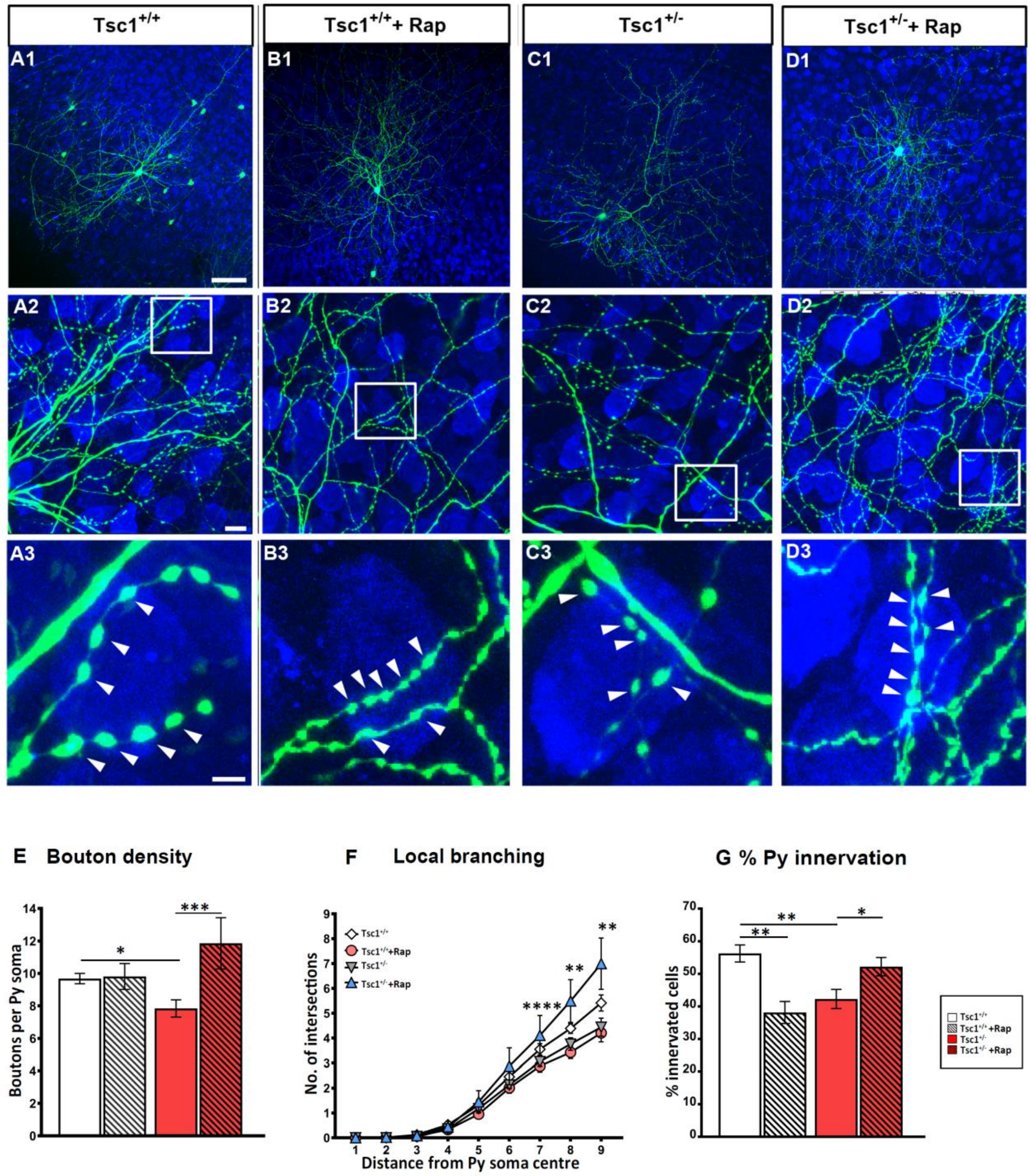


**Supplemental Figure 9. Short term Rapamycin treatment rescues loss of perisomatic innervation in Tg*(Nkx2.1-Cre);Tsc1*$^{flox/+}$ mice at EP34. A, B,** PV cell (green) among NeuN immunostained neurons (blue) in cortical organotypic cultures from *Tsc1*$^{Ctrl}$ mouse at EP34 where **B** is treated with Rapamycin. **C, D,** PV cells from Tg*(Nkx2.1-Cre);Tsc1*$^{flox/+}$ where **D** is treated with Rapamycin. ***E***, Loss of bouton density in PV cells from Tg*(Nkx2.1-Cre); Tsc1*$^{flox/+}$ mice is reversed by Rapamycin treatment (two-way ANOVA with Bonferroni post hoc analysis, *p<0.05, ***p<0.001 ); PV cells: n=14 *Tsc1*$^{Ctrl}$, n=14 Tg(*Nkx2.1-Cre*);*Tsc1*$^{flox/+}$, n=8 *Tsc1*$^{Ctrl}$ + Rapamycin, n=6 Tg(*Nkx2.1-Cre*); *Tsc1*$^{flox/+}$+ Rapamycin. Similarly, loss of local branching in PV cells from Tg(*Nkx2.1-Cre*); *Tsc1*$^{flox/+}$mice is reversed by Rapamycin treatment (**F**) (two-way ANOVA with Bonferroni post hoc analysis, **p<0.001, ***p<0.001). **G**, Rapamycin treatment reverses the loss of percentage of innervation in mutant PV cells, while it leads to significant loss of innervation in PV cells from *Tsc1*$^{Ctrl}$ mice (two-way ANOVA with Bonferroni post hoc analysis, *p<0.05, **p<0.001). Arrowheads indicate boutons. PV cells: n=12 *Tsc1*$^{Ctrl}$, n=14 Tg(*Nkx2.1-Cre*);*Tsc1*$^{flox/+}$, n=7 *Tsc1*$^{Ctrl}$ + Rapamycin, n=6 Tg(*Nkx2.1-Cre*);*Tsc1*$^{flox/+}$ + Rapamycin. Scale bars: ***A1-D1***, 10 µm; ***A2-D2*** and ***A3-D3***, 5 µm. Data represents mean ± SEM.

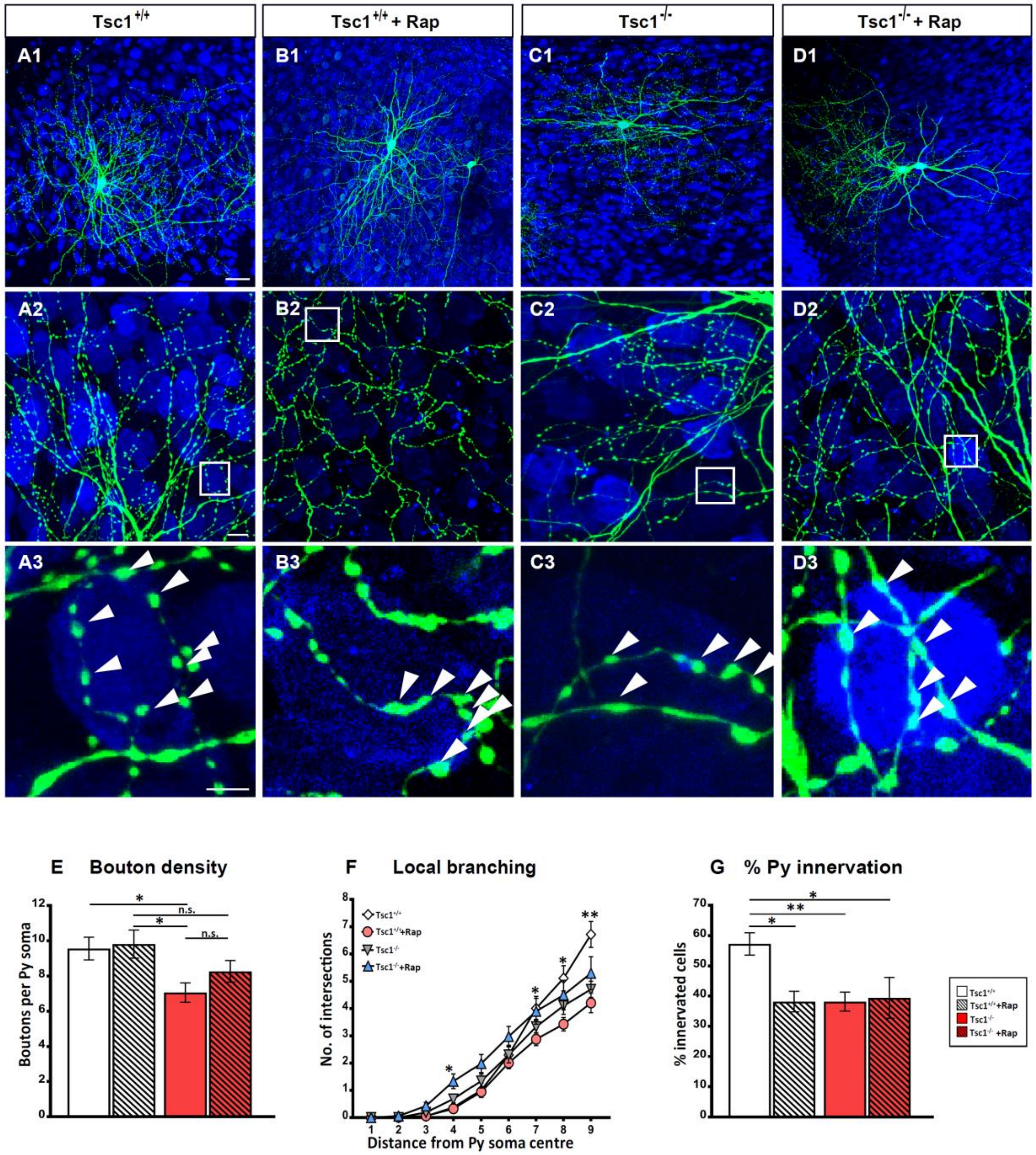


**Supplemental Figure 10. Short term Rapamycin treatment does not rescue loss of perisomatic innervations in Tg*(Nkx2.1-Cre);Tsc1*$^{flox/flox}$ mice at EP34. A, B,** PV cell (green) among NeuN immunostained neurons (blue) in cortical organotypic cultures from a *Tsc1*$^{Ctrl}$ mouse at EP34 where **B** is treated with Rapamycin. **C, D,** PV cells from

Tg(*Nkx2.1-Cre*);*Tsc1*$^{flox/flox}$ where **D** is treated with Rapamycin. **E**, **F**, Loss of bouton density (**E**) and terminal branching (**F**) in PV cells from Tg(*Nkx2.1-Cre*); *Tsc1*$^{flox/flox}$ mice are only partially reversed by Rapamycin treatment. Two-way ANOVA with Bonferroni post hoc analysis, *p<0.05, **p<0.001; PV cells: n = 14 *Tsc1*$^{Ctrl}$, n= 14 Tg(*Nkx2.1-Cre*); *Tsc1*$^{flox/flox}$, n= 8 *Tsc1*$^{Ctrl}$ + Rapamycin, n= 7 Tg(*Nkx2.1-Cre*); *Tsc1*$^{flox/flox}$ + Rapamycin. **G**, Loss of percentage of innervation in PV cells from Tg(*Nkx2.1-Cre*); *Tsc1*$^{flox/flox}$ is not reversed by Rapamycin treatment. Moreover, Rapamycin treatment in *Tsc1*$^{Ctrl}$ mice causes significant loss of innervation (two-way ANOVA with Bonferroni post hoc analysis, *p<0.05, **p<0.001). Arrowheads indicate boutons. PV cells: n=14 *Tsc1*$^{Ctrl}$, n=14 Tg(*Nkx2.1-Cre*);*Tsc1*$^{flox/flox}$, n=7 *Tsc1*$^{Ctrl}$ + Rapamycin, n=7 Tg(*Nkx2.1-Cre*); *Tsc1*$^{flox/flox}$ + Rapamycin. Scale bars: ***A1-D2***, 10 µm; and ***A3-D3***, 5 µm. Data represent mean ± SEM.

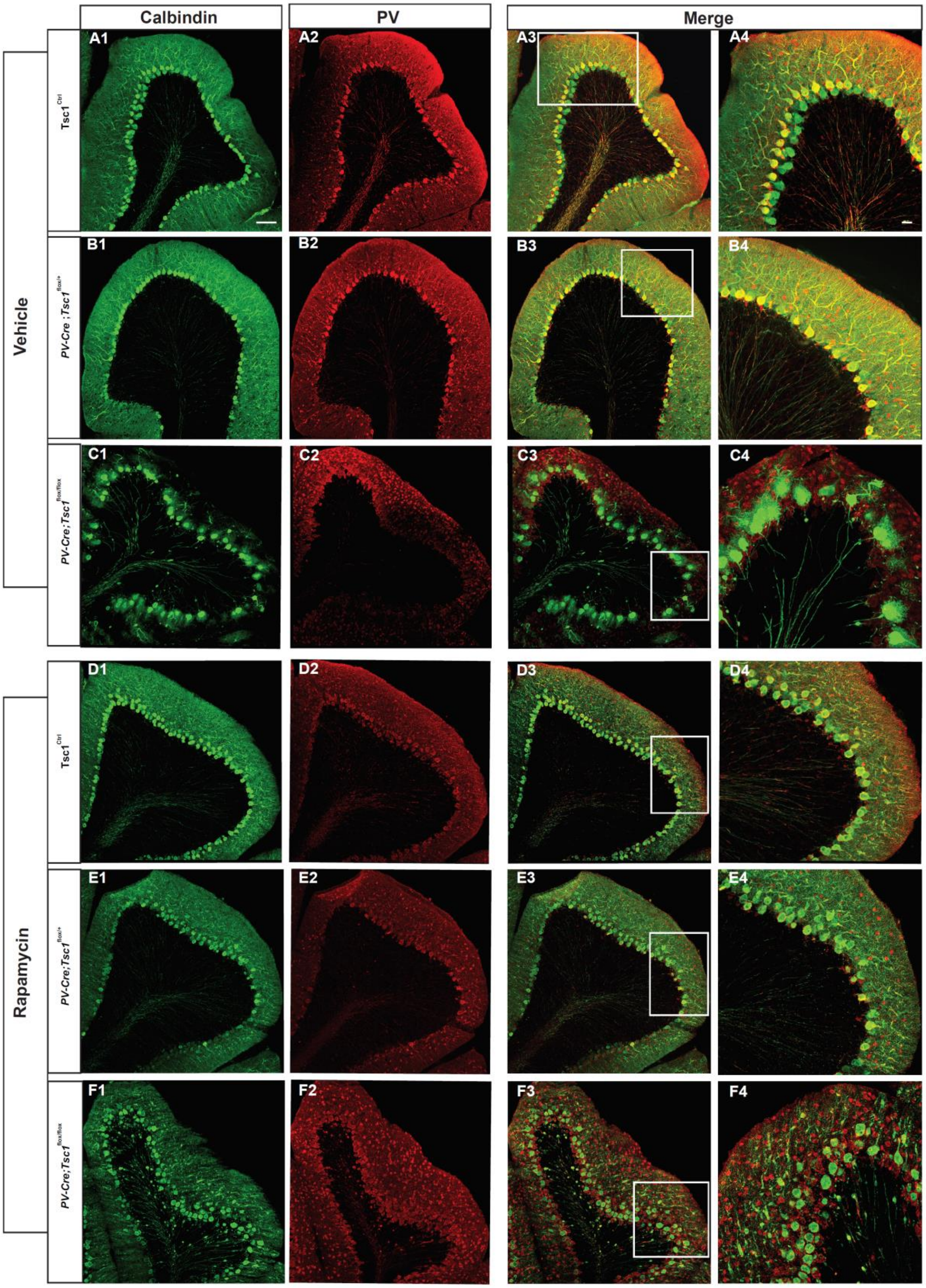

Calbindin
PV
Merge
Vehicle
Rapamycin
Tsc1^Ctrl
PV-Cre ;Tsc1^flox/+
PV-Cre;Tsc1^flox/flox
Tsc1^Ctrl
PV-Cre;Tsc1^flox/+
PV-Cre;Tsc1^flox/flox
A1
A2
A3
A4
B1
B2
B3
B4
C1
C2
C3
C4
D1
D2
D3
D4
E1
E2
E3
E4
F1
F2
F3
F4

**Supplemental Figure 11. Short term Rapamycin treatment does not rescue cerebellar abnormalities in adult homozygous mutant mice.** Coronal sections of cerebellar cortex immunostained for Calbindin (green) and PV (red) in Vehicle treated *Tsc1*$^{\mathrm{Ctrl}}$ mice (**A1-A4**), *PV-Cre;Tsc1*$^{flox/+}$ mice (**B1-B4**) , *PV-Cre*;*Tsc1*$^{flox/flox}$ mice (**C1-C4**) and Rapamycin treated *Tsc1*$^{\mathrm{Ctrl}}$ mice (**D1-D4**), *PV-Cre*;*Tsc1*$^{flox/+}$ mice (**E1-E4**), *PV-Cre*;*Tsc1*$^{flox/flox}$ mice (**F1-F4**). Note that we did not observe any obvious abnormality in *PV-Cre*;*Tsc1*$^{flox/+}$ mice, while the *PV-Cre*;*Tsc1*$^{flox/flox}$ mice were significantly affected and the rescue with rapamycin treatment was only partial (**F4**). Scale bars: ***A1-F3****,* 10 µm; and ***A4-F4***, 5 µm.
